# Chemical enhancement of DNA repair in aging

**DOI:** 10.1101/2025.02.21.639496

**Authors:** Joana C. Macedo, Maria M. da Silva, Joana M. Magalhães, Carlos Sousa-Soares, M. Inês Ala, Mafalda Galhardo, Rui Ribeiro, Monika Barroso-Vilares, Mafalda Sousa, Paula Sampaio, Elsa Logarinho

## Abstract

DNA damage is a central driver of the aging process. We previously found that KIF2C, known to play a role in DNA repair, is repressed in aged cells. Here, we investigated if increased KIF2C activity counteracts DNA damage and its effects on aging phenotypes. We show that a small-molecule agonist of KIF2C enhances DNA repair in two distinct genetic disorders exhibiting DNA damage and accelerated aging, the Hutchinson-Gilford progeria (HGPS) and Down (DS) syndromes. Mechanistically, the KIF2C agonist improves the repair of DNA double-strand breaks by inducing nuclear envelope invaginations poked by cytoplasmic microtubules, which translated into amended epigenetic and transcriptional signatures of HGPS and DS. Moreover, subcutaneous administration of the KIF2C agonist in progeria mice mitigated aging phenotypes, extending their healthspan. Our study discloses a unique geroprotective pharmacological approach targeting DNA damage.

## Main text

DNA damage affects most, if not all, aspects of the aging process, making it a potentially unifying cause of aging (*1*). Most progeroid (“premature aging-like”) syndromes are caused by mutations in genes involved in DNA repair or nuclear lamina integrity (*2*). Premature aging is also observed in long-term cancer survivors who undertook genotoxic chemo- and radiotherapy (*3*). Moreover, DNA repair proficiency appears to decline during aging (*4*). Still, therapeutic interventions directly targeting DNA damage and/or enhancing DNA repair have been largely missing.

Microtubule (MT) networks, major components of the cytoskeleton regulated by many proteins including the dynein and kinesin motors (*5*), show disrupted organization and dynamics in aged cells (*6*–*8*). Beyond their established role in chromosome segregation during mitosis (*9*), MTs have been recognized for their importance in genome maintenance and repair (*10*). The connection between cytoplasmic MTs and the linker of nucleoskeleton and cytoskeleton (LINC) complexes generates mechanical forces on the nuclear envelope that can alter nuclear shape and chromatin dynamics upon DNA damage, providing an environment conducive to DNA repair (*10*–*12*). Recent studies have supported a model in which dynamic MTs, the LINC complex, and kinesins ‘poke’ the nuclear envelope, forming nuclear envelope tubules (NETs) that facilitate double-strand breaks (DSB) reassociation and recruitment of proteins from the homologous recombination (HR) or non-homologous end joining (NHEJ) repair pathways (*13*).

We previously found found the repression of the MT-depolymerizing kinesin KIF2C in fibroblasts from elderly donors to be causally linked to chromosome mis-segregation and the accrual of aneuploid senescent cells (*14*). Genetic or pharmacological restoration of KIF2C activity in elderly cells re-established MT dynamics and chromosome segregation accuracy, thereby averting senescence. Based on the subsequently reported link between KIF2C depletion (or inhibition) and defective DSBs mobility and repair (*15*), we asked if the senostatic effect of KIF2C that we previously disclosed is due to improved enhanced DNA repair capacity. To ascertain this, we resorted to fibroblasts with increased DNA damage and nuclear deformation derived from Hutchison-Gilford Progeria Syndrome (HGPS) and Down Syndrome (DS) patients (*16, 17*), which aside the distinct genetic aetiology (*LMNA/C* mutation and chromosome 21 trisomy, respectively), exhibit features of accelerated aging in more than one organ or type of tissue (*16*).

### *KIF2C* is repressed in fibroblasts from HGPS and DS patients

To evaluate the potential genoprotective effect of a small-molecule KIF2C agonist (UMK57) in human segmental aging syndromes, HGPS and DS, we first checked if KIF2C is downregulated in cells from these patients, as previously reported for octogenarian donors (*14*). Indeed, we found significant downregulation of KIF2C protein and transcript levels in both HGPS and DS human dermal fibroblasts (HDFs) compared to age-matched healthy controls (Fig. 1A-D; Fig. S1A). We then tested if the optimal dose of UMK57 in octogenarian HDFs would also be effective in progeroid cells. Time-lapse phase-contrast microscopy of HGPS cells exposed to increasing concentrations of UMK57 confirmed that 1μM was sufficient to rescue the mitotic delay (caused by disturbed MT dynamics), while not affecting healthy controls (Fig. S1B). Also, UMK57 treatment increased the percentage of Ki67-positive proliferating cells in both HGPS and DS cultures (Fig. S1C). Moreover, attesting improved MT-depolymerizing activity, the small-molecule KIF2C agonist not only reduced the number of calcium-stable kinetochore MT fibers (k-fibers) in metaphasic HGPS and DS fibroblasts (Fig. 1E, Fig. S1D), but also reduced the rate of chromosome mis-segregation as inferred from micronuclei incidence (Fig. S1E). Importantly, to confirm that UMK57 rescues k-fiber dynamics specifically through KIF2C, we performed siRNA-mediated partial depletion of KIF2C (siKIF2C) in healthy controls and compared calcium-stable k-fibers in metaphase cells upon treatment with UMK57 vs. vehicle (Fig. S1F-H). We found that UMK57 specifically operates through KIF2C, as increased k-fiber intensity levels induced by KIF2C depletion were reduced in UMK57-treated cells. Furthermore, a chemically related analog of UMK57 differing in only one chemical group, UMK95, which does not affect MT depolymerization *in vitro* (*18*), was unable to rescue the mitotic delay and increased k-fiber intensity levels in HGPS cells (Fig. S1I, J).

**Fig. 1.**
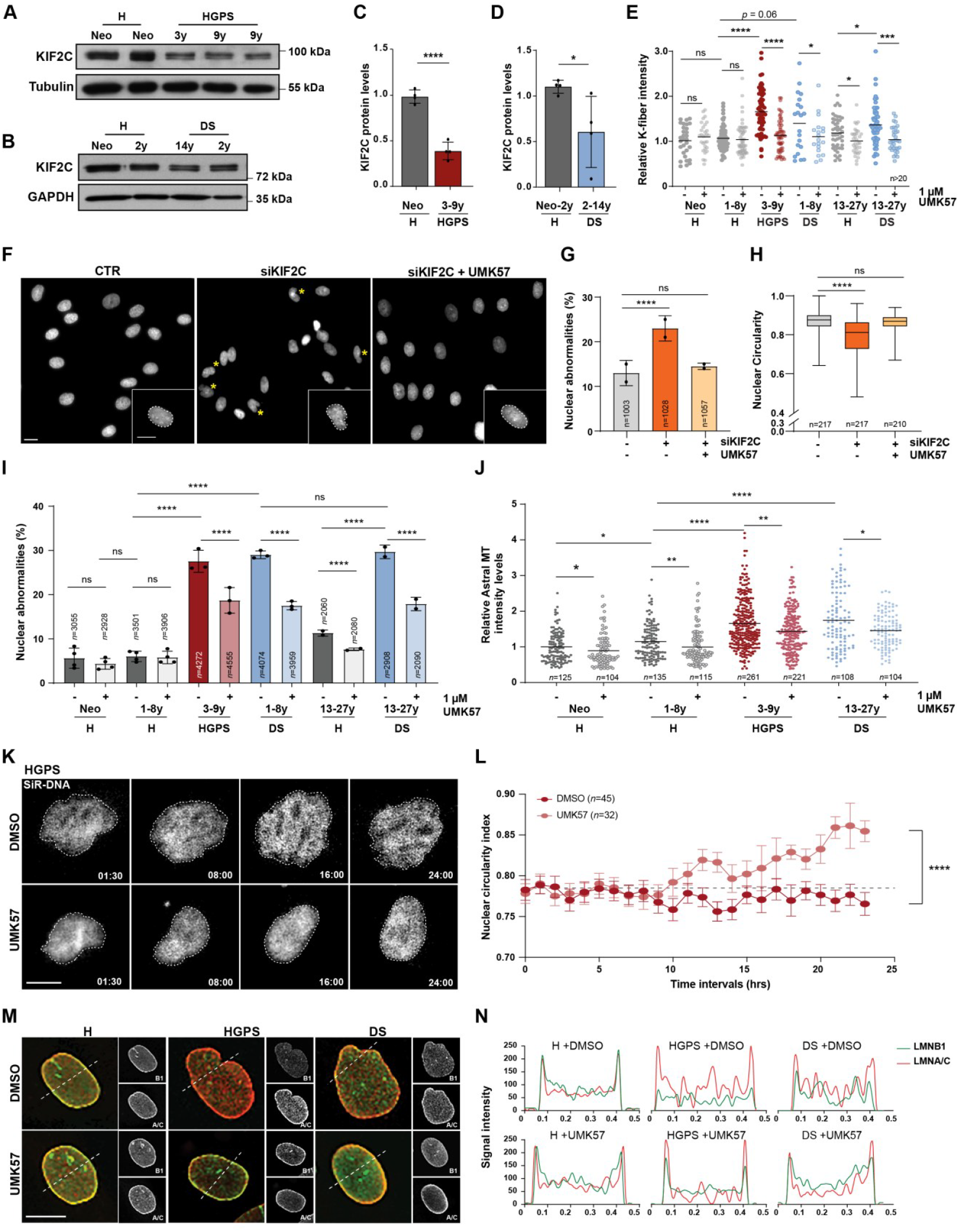
Hyperstable MTs and nuclear deformation instigated by KIF2C repression are corrected by small-molecule enhancement of KIF2C activity. **A)** Western blot analysis of KIF2C protein levels in mitotic extracts of fibroblasts from healthy (H) neonatal individuals and HGPS patients. Tubulin used as loading control. **B)** KIF2C protein levels in mitotic extracts of H and DS fibroblasts. GAPDH used as loading control. **C, D)** Western blot quantitative analyses. KIF2C protein levels were normalized to the loading controls, and H samples used as reference. **E)** Calcium-stable k-fiber intensities measured by immunofluorescence analysis of tubulin-stained metaphase cells treated with DMSO (−) and UMK57 (+) as indicated. Levels were compared to DMSO-treated neonatal cells. **F)** Representative images of nuclear morphology in neonatal cells – control (CTR), KIF2C siRNA-depleted (siKIF2C) and siKIF2C-depleted treated with UMK57 (siKIF2C+UMK57). Yellow asterisks highlight dysmorphic nuclei. **G)** Quantification of control, siKIF2C and siKIF2C+UMK57 cells with nuclear abnormalities (defined as blebs, folds, or gross irregularities in shape). **H)** Nuclear circularity indexes measured in the experimental conditions shown in F. **I)** Percentage of H, HGPS, and DS cells treated with DMSO (−) or UMK57 (+) exhibiting nuclear abnormalities. **J)** Tubulin intensity levels in astral MTs repolymerized after cold shock in cells treated with DMSO (−) or UMK57 (+). **K)** Serial images of confocal spinning-disk movies of HGPS laminopathic nuclei labeled with SiR-DNA and treated with DMSO or UMK57. **L)** Nuclear circularity indexes measured throughout live-imaging shown in K. **M)** Representative images of LMNB1 (green) and LMNA/C (red) immunostainings in H, HGPS, and DS fibroblasts treated with DMSO or UMK57. **N)** LMNB1 (green) and LMNA/C (red) immunostaining intensity profiling across the nuclear transversal sections highlighted by dashed lines in M). Scale bars, 20 μm. All values are mean ± s.d. except for H (median ± range) of at least two independent experiments. Sample size (n) is indicated in each graph. ns *p*>0.05, * *p*<0.05, ** *p*<0.01, *** *p*<0.001, **** *p*<0.0001 by unpaired t-test (C-E), Fisher’s exact (G, I), Kruskal-Wallis test followed by Dunn’s multiple comparisons test (H), Mann-Whitney (J), and one-way ANOVA (L) statistical tests.

### Hyperstable cytoplasmic MTs instigated by KIF2C repression cause nuclear deformation

Upon siKIF2C depletion, an unforeseen nuclear dysmorphology phenotype was observed in neonatal HDFs (Fig. 1F), as shown by the increased percentage of cells with nuclear abnormalities (Fig. 1G) and the decreased nuclear circularity index in the cell population (Fig. 1H). UMK57 treatment rescued this phenotype, confirming that it is driven by KIF2C repression (Fig. 1F-H). UMK57 treatment also improved nuclear shape in HGPS and DS cell cultures, which typically exhibit misshapen nuclei (Fig. 1I). In line with this, the nuclear circularity indexes of HGPS and DS cells increased with UMK57 but remained unchanged with UMK95 treatment (Fig. S2A, B). Releasing MT-generated forces on the nucleus, either by growing cells on “soft substrates” (*19*) or treating cells with the small-molecule Remodelin (*20*), was shown to improve the nuclear shape morphology and cellular fitness in laminopathic cells. Therefore, we tested if small-molecule enhancement of KIF2C activity alleviates MT-generated forces on the nuclear envelope and restores nuclear structure in HGPS and DS cells. We measured MT regrowth kinetics after cold treatment-induced MT depolymerization (Fig. S2C; t=0 min). While the initial MT nucleation (t=5 min) at the MT-organizing center appeared equally efficient in HGPS, DS and control cells, HGPS and DS cells exhibited significantly higher tubulin intensity levels at the astral MTs, indicative of hyperstability and in line with reduced KIF2C activity (Fig. 2J). UMK57 treatment reduced the density of astral MTs in HGPS and DS patients’ cells (Fig. S2C, Fig. 2J). To determine if reversion of MT hyperstability improves the nuclear shape in UMK57-treated HGPS and DS cells, we measured the nuclear circularity index of individual cells at times t=0 and t=5 min after cold shock. Nuclear circularity decreased in vehicle-treated HGPS and DS cells upon MT nucleation (t=5 vs. t=0 min) but not in UMK57-treated cells (Fig. S2D). Live-cell imaging of HGPS cells stained with a far-red SiR-DNA probe further confirmed the corrective effect of UMK57 on nuclear circularity (Fig. 2K, L). Moreover, histogram profiles of Lamin A/C and Lamin B1 immunofluorescence levels across the nuclear diameter revealed that UMK57 improves the distribution pattern towards the nuclear periphery in HGPS and DS cells (Fig. 2M, N), as well as the levels of Lamin B1 (Fig. S2E). Importantly, siKIF2C depletion resulted in impaired distribution and levels of Lamin B1, which were rescued by UMK57 treatment (Fig. S2F, G).

**Fig. 2.**
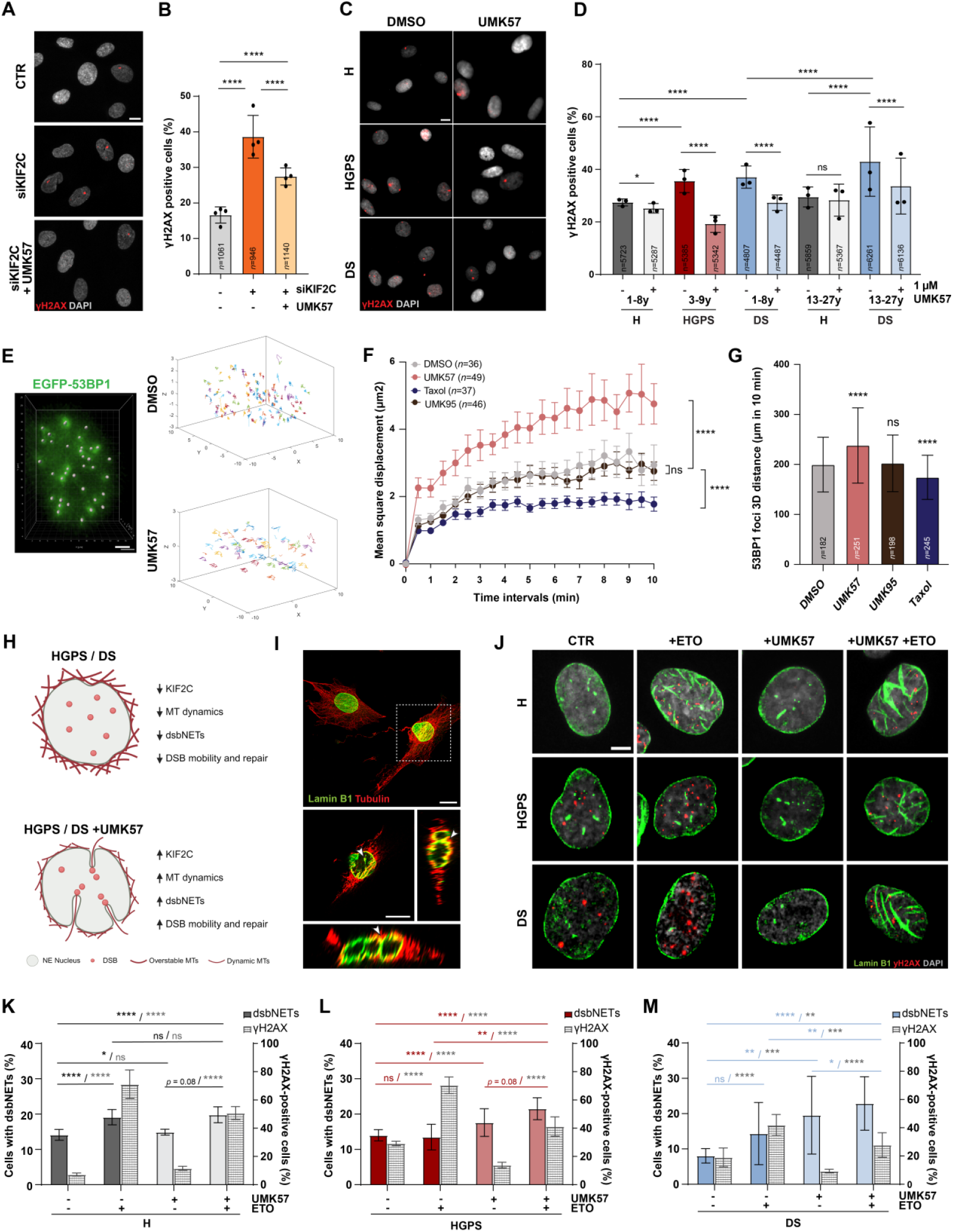
KIF2C repression induces DSB DNA damage that is rescued by improved dsbNETs-mediated DNA repair upon UMK57 treatment. **A)** Representative images of control, siKIF2C-depleted and siKIF2C+UMK57 neonatal cells immunostained for γH2AX (DSB marker; in red). **B)** Percentage of γH2AX-positive cells in the experimental conditions shown in A. **C)** Representative images of H, HGPS, and DS fibroblasts treated with DMSO (−) or UMK57 (+) and immunostained for γH2AX (in red). **D)** Percentage of γH2AX-positive cells in the experimental conditions shown in C. **E)** Examples of 10 min 3D mobility traces of EGFP-53BP1 foci in DMSO- and UMK57-treated cells under genotoxic stress induced by 1μM etoposide treatment. **F)** Measurements of mean-square displacement of EGFP-53BP1 foci in cells imaged as mentioned in E) and treated with DMSO, Taxol (5 nM; MT stabilizer), UMK95 (100 nM) or UMK57 (100 nM), as indicated. **G)** Quantification of the distance traveled by EGFP-53BP1 foci over 10 min in cells treated as described in F. **H)** Scheme illustrating the rationale behind KIF2C role in DSB DNA repair via generation of dsbNETs dependent on fitted MT dynamics. KIF2C repression in HGPS and DS cells leads to overstable MTs that are unable to promote dsbNETs in response to DSB DNA damage. UMK57 improves KIF2C activity and MT dynamics that allows for dsbNETs formation. **I)** Representative image of etoposide-treated healthy fibroblasts exhibiting dsbNETs (arrowheads), invaginating nuclear envelope tubules staining positive for LMNB1 (green) and tubulin (red) antibodies. **J)** Representative images of healthy (H), HGPS and DS fibroblasts treated with DMSO (CTR) or UMK57 for 16h, and then challenged (or not) with 10 μM etoposide (ETO) for 1h before fixation and immunostaining with LMNB1 (green) and γH2AX (red) antibodies. **K-M**) Quantitative analysis of the percentage of H (K), HGPS (L) and DS (M) cells with dsbNETs (yy axis on the left) and DSB DNA damage (yy axis on the right) in the experimental conditions mentioned in J. Scale bars: 10 μm (A, B, I), 2 μm (E) and 5 μm (J). All values are mean ± s.d. of at least three independent experiments. Sample size (n) is indicated in each graph. ns *p*>0.05, * *p*<0.05 and **** *p*<0.0001 by Fisher’s exact test (B, D, K, L, M), two-way ANOVA (F) and Mann-Whitney (G) statistical tests.

### Enhanced microtubule dynamics by the KIF2C agonist improves DNA repair

Several lines of evidence have linked changes in nuclear lamina and nuclear shape to defective DNA repair (*21, 22*). In agreement, siKIF2C-depleted laminopathic cells exhibited increased levels of unrepaired DNA damage (γH2AX DSBs marker) in comparison to controls. UMK57 treatment mitigated DNA damage in siKIF2C-depleted cells (Fig. 2A, B). Similarly, the elevated DNA damage observed in HGPS and DS cells in comparison to healthy controls was rescued upon treatment with UMK57 (Fig. 2C, D). Moreover, when challenged with the DNA damage-inducing agent etoposide, HGPS and DS cells exhibited higher levels of γH2AX in comparison to controls, indicating impaired DNA repair (Fig. S3A-C). However, concomitant treatment with etoposide and UMK57 was able to prevent the accumulation of DNA damage (Fig. S3A-C). Since KIF2C MT-depolymerizing activity was previously shown as required for the mobility and repair of DSBs (*15*), we used a live-cell reporter to ascertain if the UMK57 agonist acts to improve the mobility of etoposide-induced DSBs labeled with EGFP-53BP1 (*11*). The 3D trajectories of randomly selected EGFP-53BP1 foci (Fig. 2E) were used to calculate the mean square displacement (MSD, deviation from reference position over time) (Fig. 2F) and the total 3D distance traveled during 10 min (Fig. 2G). Chemical enhancement of KIF2C motor activity with UMK57 resulted in augmented DSBs mobility in comparison to treatment with vehicle only (Fig. 2E-G). Confirming the requirement for dynamic MTs, treatment with the MT stabilizing agent taxol diminished both MSD and 3D distance parameters (Fig. 2F, G). Demonstrating that UMK57 acts specifically via KIF2C, the UMK95 inactive drug analog showed no effect in the DSBs dynamics (Fig. 2F, G), nor in the mitigation of basal or etoposide-induced DNA damage in HGPS cells (Fig. S3D-F).

Recent studies have uncovered a nuclear-cytoplasmic DNA damage response mechanism in which several molecular players collaborate to generate DSB-capturing nuclear envelope tubules (dsbNETs) that facilitate DNA repair (*11, 13*). We asked if KIF2C improves DSBs mobility and repair by facilitating the assembly of dsbNETs in HGPS and DS cells with disrupted MT dynamics (Fig. 2H). Immunofluorescence analysis of MTs and nuclear lamina was used to inspect the formation of dsbNETs in healthy fibroblasts treated with etoposide. DSB DNA damage induced the assembly of nuclear tubules infiltrated by MTs as perceived by the double-positive staining for Lamin B1 and tubulin (Fig. 2I). Next, we quantified the percentage of cells exhibiting dsbNETs in healthy, HGPS and DS cultures treated with vehicle or UMK57 for 16h before exposure to etoposide for 1h (Fig. 2J). Here, double immunostaining of Lamin B1 and γH2AX was used to detect dsbNETs and DSBs, respectively. We found that vehicle-treated HGPS and DS cells inefficiently assembled dsbNETs in comparison to healthy controls despite the higher levels of γH2AX^+^ DSBs induced by etoposide. Notably, the percentage of HGPS and DS cells with dsbNETs increased upon UMK57 treatment, concomitantly with a reduction in γH2AX^+^ cells, indicating that enhanced KIF2C activity restores the MT dynamics required for dsbNETs assembly and DSBs repair in response to etoposide (Fig. 2J-M). Again, siRNA-mediated depletion of KIF2C in healthy cells was used to confirm that the defective dsbNETs phenotype is driven by low KIF2C activity and specifically rescued by UMK57 (Fig. S3G, H).

### UMK57 amends epigenetic and transcriptomic signatures of aging in HGPS and DS cells

DNA damage and nuclear envelope defects lead to a global loss of heterochromatin regions and changes in their histone marks (*23, 24*). Since the KIF2C agonist compound efficiently improved DNA repair and the nuclear lamina in HGPS and DS cells, we asked if epigenetic shifts in heterochromatin-associated histones could be also rescued by UMK57. Quantitative immunofluorescence analysis of the trimethylated histones H3K9me3 and H4K20me3, reported as typically down- and upregulated in HGPS cells respectively (*25*), revealed that UMK57 reverts the intensity levels in comparison to vehicle in both HGPS and DS cells (Fig. 3A and Fig. S4A).

**Fig. 3.**
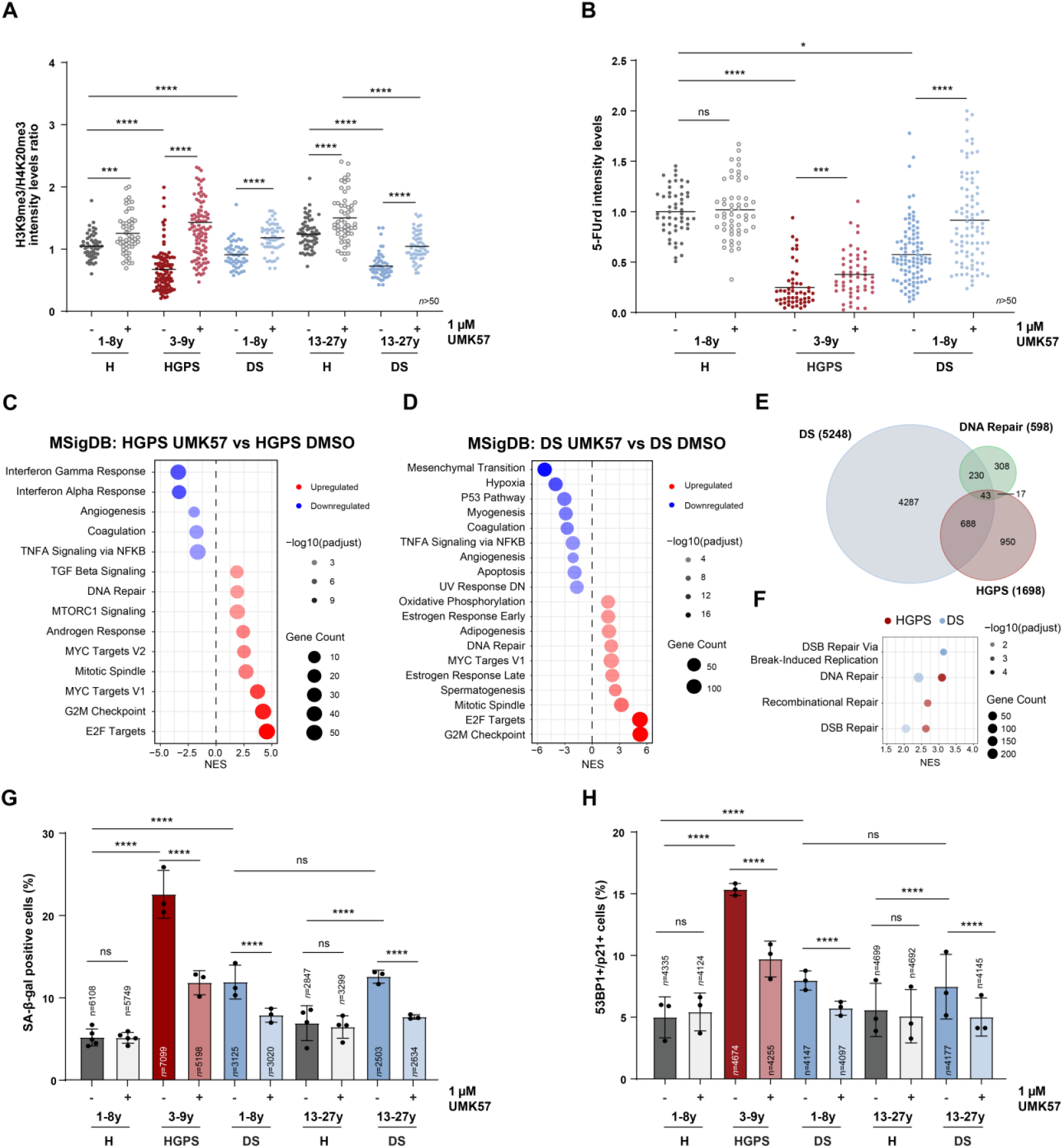
UMK57 treatment resets heterochromatin epigenetic marks and the transcriptome of HGPS and DS fibroblasts. **A)** Ratio between fluorescence intensity levels of H3K9me3 and H4K20me3 heterochromatin repressive marks in fibroblasts from H, HGPS and DS donors treated with DMSO (−) and UMK57 (+). DMSO-treated neonatal sample was used as reference. **B)** Intensity levels of fluorescent 5-FUrd incorporated into RNA transcripts in H, HGPS and DS donors treated with DMSO or UMK57. DMSO-treated neonatal sample was used as reference. **C, D)** Gene set enrichment analysis (GSEA) for HGPS (C) and DS (D) fibroblasts treated with UMK57 vs. DMSO using the hallmark gene sets from the MSigDB database. Significantly altered terms were plotted accordingly to the ascending normalized enrichment score (NES) (blue, negative score; red, positive score). **E)** Venn diagram showing the overlap between the genes found differentially expressed in DS and HGPS fibroblasts upon UMK57 treatment, and DNA repair genes from MSigDB (GO:0006281). **F)** GSEA of DNA repair pathways from MSigDB (biological processes) significantly modulated by UMK57 treatment in HGPS (red) and DS (blue). Significantly altered terms were plotted accordingly to NES descending order. **G)** Percentage of H, HGPS and DS cells treated with DMSO (−) or UMK57 (+) with detectable SA-β-galactosidase (SA-β-gal) activity. **H)** Percentage of H, HGPS and DS cells treated with DMSO (−) or UMK57 (+) staining positive for both the cell cycle inhibitor CDKN1A/p21 and the DNA damage 53BP1 senescence markers. All values are mean ± s.d. of at least three independent experiments. Sample size (n) is indicated in each graph. ns *p*>0.05, * *p*<0.05 and **** *p*<0.0001 by Mann-Whitney (A, B) and Fisher’s exact test (G, H) statistical tests. Significance scores of functional enrichment analyses (C, D, F) are described in Methods.

To ascertain for the impact of a potentially improved chromatin architecture in gene expression, we measured the transcription activity and characterized the transcriptome of HGPS and DS treated with UMK57 vs. vehicle. Immunolabelling of nascent RNAs with 5-FUrd showed that active transcription is reduced in HGPS and DS cells in comparison to healthy controls (*26*), but enhanced upon UMK57 treatment (Fig. 3B). RNA-sequencing datasets were generated from samples of HGPS and DS fibroblasts treated with vehicle or UMK57. Although the number of differentially expressed genes (DEGs) that were downregulated by the UMK57 treatment was similar in both HGPS and DS cells, the number of upregulated DEGs was extensively higher in DS cells (Fig. S5A, B). However, gene-set enrichment analysis (GSEA) for pathways from the Molecular Signatures Database (MSigDB) revealed that the upregulated pathways driven by UMK57 treatment are mostly common between HGPS and DS cells, including cell cycle signatures associated with E2F and MYC transcriptional activity, mitosis and DNA repair, overall supporting an improved cell fitness (Fig. 3C, D). Hierarchical clustering of enriched GO terms for common upregulated genes in UMK57-treated HGPS and DS cells further underlined the upregulation of cell cycle, cytoskeleton assembly, chromosome segregation and DNA replication, repair and conformation biological processes (Fig. S5C). GSEA for 43 genes of the DNA repair MSigDB (GO:BP) that were found to be differentially expressed in both HGPS and DS conditions (Fig. 3E and Fig. S5A-D) revealed that UMK57 primarily boosts DSB repair pathways (Fig. 3F), in agreement with the amended DSB mobility and dsbNETs assembly phenotypes mentioned above. Also, overrepresentation analysis shows an enrichment in several DNA repair processes for commonly upregulated genes in HGPS and DS following UMK57 treatment, in agreement with common DNA repair DEGs being mostly upregulated in both conditions (Fig. S5D, E). Upregulation of genes with functional roles in DSB DNA repair was additionally confirmed by RT-qPCR analysis (Fig. S5F) and immunofluorescence (Fig. S5G, H). Regarding the downregulated molecular signatures induced by the UMK57 treatment, these mainly included pro-inflammatory pathways in HGPS (e.g. IFN-γ and IFN-α responses, TNF-α signaling via NF-κB), and DNA damage response pathways in DS (e.g. p53 pathway, UV response, TNF-α signaling via NF-κB) (Fig. 4C). In agreement with an improved transcriptome profile, we found reduced percentage of HGPS and DS cells exhibiting senescence markers, namely senescence-associated β-galactosidase activity (Fig. 4G), and DNA damage (53BP1)/cell cycle inhibition (CDKN1A/p21) (Fig. 4H). Moreover, the senescence phenotype depends on the activation of the cGAS-STING pathway of the innate immune response, which links genomic instability to inflammation (*27*), since siRNA-mediated depletion of cGAS (Fig. S6A-D) rescued the higher percentage of senescent cells in HGPS and DS cultures (Fig. S6F, G). Overall, our findings demonstrate that chemical enhancement of KIF2C motor activity boosts DNA repair via improved dsbNETs assembly, restoring nuclear architecture and chromatin organization towards a transcriptional landscape preventive of inflammatory gene expression and senescence.

**Fig. 4.**
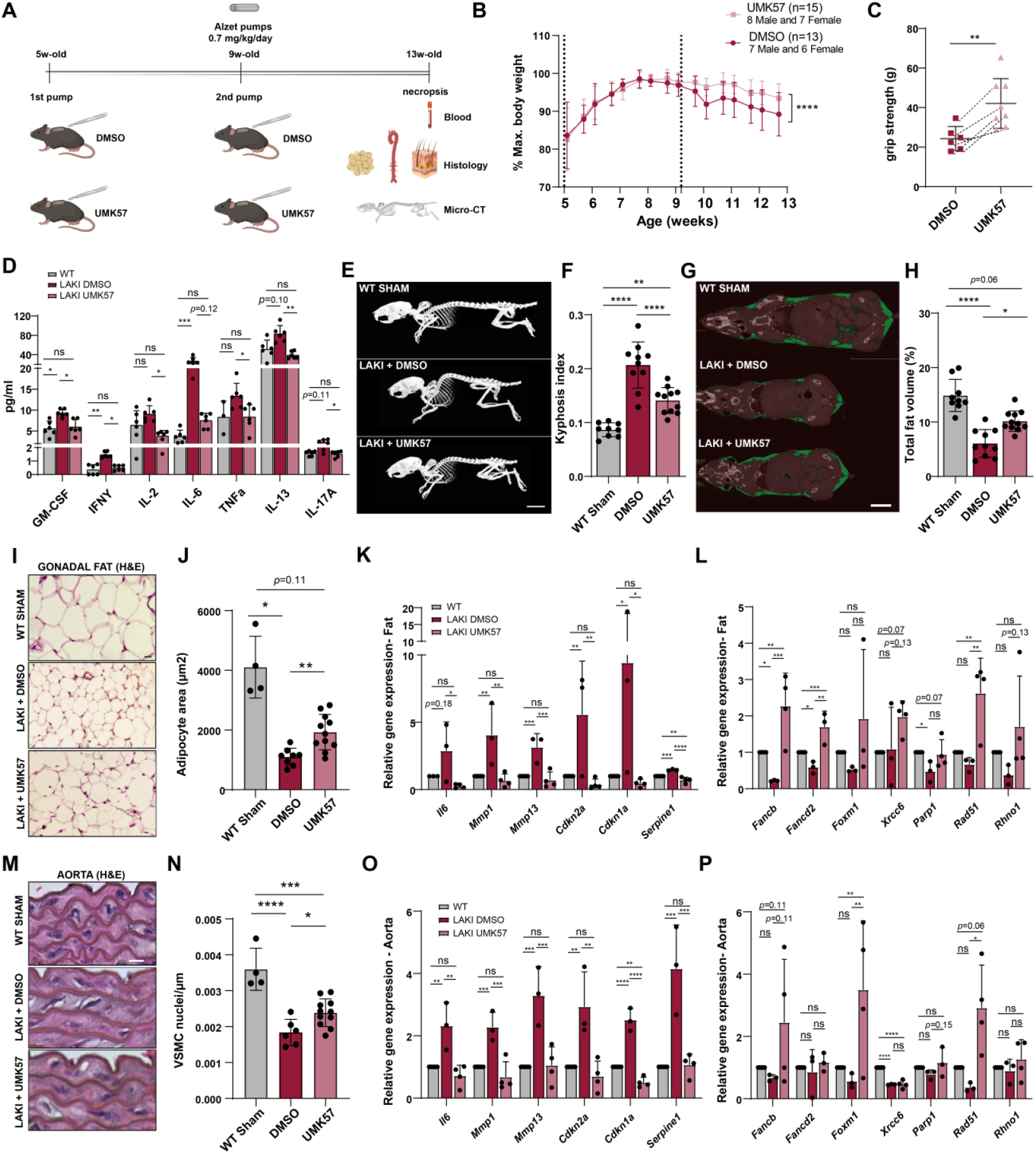
Small-molecule Kif2C induction *in vivo* delays histopathologic features in progeria mice. **A)** Experimental layout of the pre-clinical validation of the UMK57 treatment in progeria mice (*Lmna*^*G609G/G609G*^ or LAKI). UMK57 (or DMSO control) was delivered via serial subcutaneous implants of two osmotic pumps. 13-week-old animals were then used for phenotypic analyses. **B)** Body weight analysis in LAKI mice treated with DMSO or UMK57 for 8 weeks. **C)** Grip-strength test in 11-week-old LAKI mice treated with DMSO or UMK57. Dashed lines are used to highlight littermates. **D)** Cytokine levels in the serum of WT (reference) and LAKI mice treated with DMSO or UMK57. **E)** Representative 3D reconstructions of micro-CT segmented images of the whole-skeleton. WT mice subjected to pump implants (sham control) were used as reference. **F)** Kyphosis index based on skeleton 3D reconstructions. **G)** Representative 3D reconstructions of micro-CT segmented images for whole-body fat (green). **H)** Total fat volume based on 3D reconstructions. **I)** Histological analysis of gonadal fat and **J)** quantification of adipocyte cross-sectional area (>50 adipocytes per animal). **K, L**) Expression levels of senescence-associated (K) and DNA repair (L) genes in gonadal fat measured by RT–qPCR (2^−ΔΔCt^). **M)** Histological analysis of the aorta and **N)** VSMC nuclei density in the medial wall. **O, P**) Expression levels of senescence-associated (O) and DNA repair (P) genes in the aorta measured by RT–qPCR (2^−ΔΔCt^). Scale bars: 1 cm (E, G), 50 μm (I) and 10 μm (M). In graphs, dots represent number of animals used and values are mean ± s.d. ns *p*>0.05, * *p*<0.05 and **** *p*<0.0001 by two-way ANOVA (B), unpaired t-test (C), Kruskal-Wallis followed by Dunn’s multiple comparisons (F, H, J, N) ordinary one-way ANOVA parametric test with Tukey’s multiple-comparison correction (K, L, O, P) statistical tests.

### Chemical enhancement of KIF2C activity mitigates histopathological features of progeria mice

We next investigated the geroprotective effect of the UMK57 pharmacological intervention in a HGPS mouse model (*Lmna*^*G609G/G609G*^ or LAKI mice) (*28*). We started by testing a short-term treatment in adult mouse fibroblasts (MAFs) isolated from the ear dermis of LAKI mice. Decreased incidence of misshapen nuclei and micronuclei in LAKI MAFs were used as readouts to determine the optimal dose of UMK57 capable of restoring MT dynamics while having no impact in WT controls (i.e. 0.5μM) (Fig. S7A, B). LAKI MAFs treated with UMK57 for 4 days exhibited improved proliferative capacity (Fig. S7C) and decreased DNA damage (Fig. S7D), senescence (Fig. S7E) and nuclear abnormalities (Fig. S7F) in comparison to vehicle-treated controls. We additionally tested a cyclic scheme, consisting of 2 days treatment followed by 3 days withdrawal and again 2 days treatment, to monitor the perdurance of improved cellular phenotypes. We observed sustained reduction of faulty phenotypes in the LAKI MAFs under cyclic UMK57 treatment (Fig. S7G-J).

To evaluate the drug suitability for *in vivo* studies, we first assessed the thermodynamic solubility, the rate of flux across polarized Caco-2 cell monolayers, and the clearance by hepatic CYP-mediated metabolism and metabolites formed (Tables S1-S3). Although the Caco-2 permeability assay for intestinal absorption was encouraging, the low solubility (<0.025 mg/mL) and fast CYP-mediated metabolism (1.5 min half-time) of the UMK57 compound precluded the use of oral and intravenous administration routes. Therefore, we pursued with subcutaneous administration via implantation of osmotic pumps, which allowed controlled drug delivery at a rate of 0.7 mg/kg/day for 4 weeks. We followed an experimental timeline of a 2-month treatment period, with the first pump implanted in 5-week-old animals and replaced by a new pump upon 4 weeks (Fig. 4A). LAKI mice were treated with either vehicle or UMK57 and compared to sham WT controls. Micro-computed tomography (micro-CT) and necropsies were performed at the experimental endpoint. Blood samples and tissues (gonadal fat, aorta, skin, spleen, kidney and liver) were collected for biodistribution, histology and qRT-PCR analyses (Fig. 4A). Mass spectrometry analysis confirmed the presence of UMK57 in several tissues (Fig. S8A). Body weight assessment revealed that the typical weight loss starting in 9-week-old LAKI animals was deferred by the UMK57 treatment (Fig. 4B), alongside with an improvement of the forelimbs muscles as measured by the grip strength test (Fig. 4C). Moreover, UMK57-treated animals exhibited reduced levels of pro-inflammatory cytokines in the blood serum (Fig. 4D). micro-CT analysis showed that the kyphosis index (spine curvature) and lipodystrophy (reduced body fat) progeroid features typically present in LAKI mice were mitigated in UMK57-treated mice (Fig. 4E-H). Tibia micro-CT imaging displayed also substantial improvements in the bone microarchitecture of LAKI mice treated with UMK57 vs. vehicle, namely increased bone mineral density (BMD), trabecular bone volume fraction (BV/TV), trabecular number (Tb.N), and trabecular separation (Tb.Sp) (Fig. S8G-K). Histological analysis of the gonadal fat further disclosed a marked enlargement of the average adipocyte cross-sectional area in the LAKI mice treated with UMK57 (Fig. 4I, J), concurrently with a downregulation of senescence-associated genes (Fig. 4K) and an upregulation of DNA repair genes (Fig. 4L) as previously observed *in vitro*. In addition, the aortic wall thickening and the loss of vascular smooth muscle cells (VSMCs) in the middle aortic layer of LAKI animals were rescued by the drug treatment (Fig. 4M, N and Fig. S8B), and again along with improved gene expression profiles of senescence and DNA repair pathways. Histological analysis of the dorsal skin also showed that the UMK57 treatment mitigates the atrophy of the subcutaneous fat layer and thinning of the epidermal layer in LAKI mice (Fig. S8C, D) while inhibiting senescence markers (Fig. S8E) and enhancing DNA repair gene expression (Fig. S8F). Altogether, we provide pre-clinical evidence for the geroprotective role of an unprecedented chemical boosting of DNA repair via the upregulation of the KIF2C MT-depolymerizing activity.

## Conclusions

Accumulating evidence points to DNA damage as primary driver of aging (*1*). DNA damage leads to epigenetic erosion (*29*) and changes in 3D genome architecture that instigate loss of cell identity and function and ultimately trigger the senescence state (*30*). Moreover, most premature aging-like syndromes are caused by mutations in genes essential for genome stability (*31*), and longer-lived species share higher expression of core genes involved in multiple DNA repair pathways (*32, 33*). However, interventions targeted at restoring genomic integrity by augmenting DNA repair are still missing, mainly due to the fact that repair of DNA lesions requires the activities of many multi-enzyme pathways that cannot be simply enhanced by the upregulation of one or a few genes. This study demonstrates that chemical correction of the defective KIF2C motor activity in progeroid cells, mitigates DNA damage by improving genome stability mechanisms that require adequate levels of MT-depolymerizing activity. Specifically, KIF2C activation enhances nuclear-cytoplasmic DNA damage response, a process in which nuclear envelope tubules guided by dynamic MTs poke the nucleus to facilitate chromatin mobility-dependent repair (*13*). Solid proof has pointed to MT dynamics crucial function in genome maintenance (*10, 34*), with several MT plus-ended-directed kinesins reported to play a role in DSB repair (*11, 13, 15*). Among these, KIF2C holds the advantage of being targetable by the UMK57 drug agonist, excitingly conveying a pharmacological intervention for age-associated DNA damage. Notably, correction of DNA damage was shown to occur concomitantly with improved nuclear shape and chromatin reorganization, restored gene expression of cell cycle-related pathways, including DNA repair, and repression of senescence-associated inflammatory pathways, reinforcing DNA repair as main determinant of aging (*35*). Furthermore, the genoprotective effect of the UMK57 compound was validated in fibroblasts from patients with HGPS and DS syndromes, both of which exhibit low KIF2C levels and defective dsbNETs assembly. Whereas HGPS is the most well-characterized progeroid laminopathy in which a mutation in the *LMNA* gene has been causally linked to genomic instability, DS just recently gained evidence for a systemic progeroid status caused by increased DNA damage due to dose imbalance of the chromosome 21 gene *DYRK1A* (*17*). Therefore, this work offers a new therapeutic avenue for these and perhaps other premature aging disorders. Pre-clinical validation was conducted using progeria mice treated with UMK57. Although turning the drug suitable for oral or intravenous administration routes will demand further efforts, subcutaneous delivery via osmotic pumps was sufficient to mitigate the major progeroid features, i.e. lipodystrophy, kyphosis, osteoporosis, and aortic thickening (*36*). Importantly, the UMK57 treatment enhanced DNA repair and inhibited senescence gene expression in mechanosensitive adipose and aortic tissues (*37*). The cardiovascular benefits are of particular clinical relevance, as advanced coronary disease is leading the cause of death in HGPS patients. In comparison to other drug treatments reported to ameliorate HGPS defects - Lonafarnib, a farnesyl transferase inhibitor, and Remodelin, an N-acetyltransferase 10 inhibitor, both of which impact many cellular targets, UMK57 mechanism of action was confirmed to be specifically through the KIF2C target and to reset the transcriptome towards improved cellular fitness. This reduces the risk of side effects besides offering a unified intervention against phenotypes that other drugs leave unaffected (*38, 39*). Pre-clinical validation of the UMK57 treatment in the DS mouse model should also follow, considering its auspicious senotherapeutic value to tackle DS features of premature aging (*40*), in particular immunosenescence, where DNA damage appears to be a central driver (*41*). Finally, as we previously found KIF2C to be repressed also in human octogenarian fibroblasts (*14*), we anticipate that UMK57 might offer therapeutic opportunities in broader settings of age-associated diseases, for instance in cognitive decline, as KIF2C was shown as required for synaptic plasticity and cognition in mice (*42*).

## Acknowledgments

We thank Benjamin H Kwok (Institute for Research in Immunology and Cancer - IRIC), Université de Montréal, Canada) for valuable reagents. We thank the personnel at i3S Scientific Platforms for technical support: Animal Facility (S. Lamas), CCGEN (P. Magalhães), Bioimaging (M. Lázaro), Genomics (A.M. Rocha), Advanced Light Microscopy (P. Sampaio), and Biosciences Screening (A. Maia). We are grateful to all lab members for their helpful discussions.

## Author contributions

Conceptualization: EL, JCM

Methodology: JCM, MMS, JMM, CSS, MIA, RR, MB-V, PS, EL

Software: CSS, MG, MS

Investigation: All authors Visualization: JCM, MMS, CSS, EL

Funding acquisition: EL

Project administration: EL

Supervision: EL, JCM

Writing – original draft: JCM, MMS, CSS, EL

## Data and materials availability

RNA sequencing data have been deposited in the XXX under the accession number XXX. Data associated with this study are available in the article. Any requests for data or material should be directed to, and will be fulfilled by, the senior corresponding author E. L. Further information is available in the XXX linked to this article.

## Supplementary Materials

Materials and Methods Supplementary Text

Figs. S1 to S8 Tables S1 to S6

