## Supplemental Files (Methods, Figures, Tables) for "Chemical enhancement of DNA repair in aging"

### Materials and Methods

Cell culture

All human dermal fibroblasts (HDFs) used in this study are described in Table S4. Primary cell cultures were grown in Minimal Essential Medium (MEM; 10-010-CV, Corning) supplemented with 15% fetal bovine serum (FBS; 10270106, Gibco) and antibiotic–antimycotic (AA; 15240062, Gibco), and used at early passages (p≤4) as previously reported (*43*). HeLa cell line (ATCC®) was cultured in DMEM high-glucose (11330032, Gibco) with 10% FBS (10270106, Gibco) and 1x AA (15240062, Gibco). Mouse adult fibroblasts (MAFs) were isolated from ear biopsies. Biopsies were cut into small pieces and incubated with 3 mg/mL collagenase type II (17101-015, Gibco) in Dulbecco's modified Eagle's medium supplemented with F12 nutrient mixture (DMEM:F12, 11330032, Gibco) for 45 min at 37 °C in 5% $CO^2$. MAFs were then allowed to grow in a 6-well plate containing DMEM:F12 supplemented with 10% FBS (10270106, Gibco) and 1x AA (15240062, Gibco). Only early passage cultures (p<3) were used in all experiments.

Drug treatments

For fixed-cell experiments, cells were incubated for 48 h with 100 nM (HeLa cells), 1 µM (HDFs) and 0.5 µM (MAFs) of UMK57 (#342595-74-8, AOBIOUS) or UMK95 (kindly provided by Dr. Benjamin Kwok). For live-cell imaging, UMK57 or UMK95 were added to cell medium prior to starting imaging. Proteasome inhibitor MG-132 (474790, EMD Millipore) was used at 5µM for 2h. S-trityl-L-cysteine (STLC)/Eg5 inhibitor (2799-07-7, Tocris) was used at 5µM for 16h. For mitotic enrichment, asynchronous cell cultures were incubated for 16h with 5 µM of S-trityl-L-cysteine (STLC)/Eg5 inhibitor (2799-07-7, Tocris). DNA damaging agent etoposide (S1225, Selleck Chemicals) was used at 0.1 µM for 48 h in western blot analyses, 1 µM overnight in live-cell imaging, and 5-10 µM for 1 h to evaluate dsbNETs. Taxol (T7402, Sigma-Aldrich) was used at 0.5 µM to stabilize microtubules.

KIF2C siRNA knockdown

Cells were plated in serum-free culture medium and transfected 1 h later with siRNA oligonucleotides specifically targeting *KIF2C* (Merck, 5'-GAUCCAACGCAGUAAUGGU-3') at a final concentration of 15 nM. Transfections were performed using Lipofectamine RNAiMAX in Opti-MEM medium (both from Thermo Fisher Scientific) according to manufacturer's instructions. The transfection medium was replaced 6 h later by complete medium. All experiments were performed 72 h post-transfection, and protein depletion was confirmed by Western blot.

Plasmid transfection

pEGFP-h53BP1 plasmid (a gift from Chris Kok-Lung Chan; Addgene #110301) was transfected into HeLa cells using Lipofectamine 2000 (Invitrogen), according to the manufacturer's instructions.

Calcium-stable k-fiber analysis

Fibroblasts grown on sterilized fibronectin-coated glass coverslips were incubated in calcium buffer (100 mM PIPES, 1 mM MgCl2, 1 mM CaCl2, 0.5% Triton X-100, pH = 6.8) for 5 min and fixed with 4% paraformaldehyde + 0.25% glutaraldehyde in PBS for 15 min, both at 37°C. Cells were then rinsed first in PBS, then in TBS (50 mM Tris–HCl, pH = 7.4, 150 mM NaCl), and permeabilized in TBS + 0.3% Triton X-100 for 7 min. Blocking was performed with TBS-T (TBS + 0.05% Tween-20) + 10% FBS for 1 h, followed by incubation overnight at 4ºC with mouse anti-α-Tubulin antibody (T5168, Sigma-Aldrich, 1:1,500) in TBS-T + 5% FBS. Alexa Fluor® 568 secondary antibody (Life Technologies, USA) was used at 1:1,500 and nuclei were counterstained with DAPI (Sigma-Aldrich).

Immunofluorescence

Cells were cultured on ibiTreat μ-Plate 24-well (82406, ibidi GmbH) or sterilized glass coverslips coated with 50 μg/mL fibronectin (F1141, Sigma-Aldrich) and fixed with either 4% paraformaldehyde for 20 min at room temperature (RT) (for immunostainings with anti-Ki67, anti-γH2AX, anti-53BP1, anti-p21, anti-LaminA/C, anti-LaminB1 and anti-α-Tubulin antibodies), or with methanol for 5 min at -20 ºC (for immunostainings with anti-H3K9me3 and anti-H4K20me3 antibodies). Afterwards, cells were rinsed with phosphate-buffered saline (PBS) and then permeabilized with PBS + 0.3% Triton X-100 (0.5% in dsbNETs analysis) for 7-10 min at RT. Cells were blocked with 10% FBS in PBS-T (PBS + 0.05% Tween-20) for 1 h, followed by incubation overnight at 4ºC with primary antibodies diluted in PBS-T + 5% FBS as follows: rabbit anti-53BP1 (4937, Cell Signaling Technology, 1:100), mouse anti-p21 (SC-6246, Santa Cruz Biotechnology, 1:800), rabbit anti-Ki67 (ab15580, Abcam, 1:1,200), mouse anti-γH2AX (05 636, Sigma-Aldrich, 1:1,000), rabbit anti-H3K9me3 (ab8898, Abcam, 1:2,000), mouse anti-H4K20me3 (SC 134216, Santa Cruz Biotechnology, 1:250), mouse anti-LaminA/C (#4777, Cell Signaling Technology, 1:100), rabbit anti-LaminB1 (AB16048, Abcam, 1:250) and mouse anti-α-tubulin (T5168 Sigma-Aldrich, 1:1,500). Upon rinse in PBS-T for 3 x 5 min, cells were incubated for 1 h at RT with secondary antibodies diluted in PBS-T + 5% FBS as follows: Alexa Fluor® 488 (A11008, ThermoFisher Scientific, 1:1,500) and Alexa Fluor® 568 (A11031, ThermoFisher scientific, 1:1,500). Cells were then rinsed twice with PBS-T for 5 min and counterstained with DAPI (1:5,000 dilution; Merck-Aldrich) for 10 min at RT. Coverslips were mounted on slides (Epredia) using a homemade mounting solution (90% glycerol, 0.5% N-propyl-gallate, 20 nM Tris, pH 8).

Microtubules regrowth assay

Microtubules were depolymerized by incubation with cold medium (4 °C) on ice for 10 min. Cold medium was replaced by pre-warmed (37 ºC) culture medium, supplemented with either DMSO or UMK57, and left incubating for 5 min at 37°C to allow for microtubule regrowth. Cells were fixed with 4% paraformaldehyde + 0.25% glutaraldehyde in PBS for 15 min at 37°C, and incubated with mouse anti-α-Tubulin antibody (T5168, Sigma-Aldrich, 1:1,500) as described above.

Nascent RNA immunolabeling

Cells grown on coverslips were incubated for 20 min with 2 mM 5-fluorouridine (5-FUrd; F5130, Sigma-Aldrich), then fixed with 1% paraformaldehyde for 20 min, and permeabilized with 0.5% Triton X-100 for 7 min. Immunostaining with anti-BrdU antibody (Thermo Fisher Scientific) and with AlexaFluor® 488 secondary antibody (1:1,500; Life Technologies) followed.

Senescence associated-β-gal (SA-β-gal) assay

Cells were incubated for 90 min in medium supplemented with 100 nM Bafilomycin A1 (B1793, Sigma-Aldrich) to induce lysosomal alkalinization. The fluorogenic substrate for β-galactosidase, fluorescein di-β-D-glucopyranoside (33 μM; F2756, Sigma-Aldrich), was then added, and incubation done for another 90 min. Cells were fixed in 4% paraformaldehyde for 15 min and permeabilized with 0.1% Triton X-100 in PBS for 15 min. Nuclei were counterstained with DAPI (Sigma-Aldrich).

Fixed-cell microscopy analyses

For the analyses of calcium-stable k-fiber intensities and levels of KU80, 5-FUrd and H3K9me3 and H4K20me3, images were obtained using a Zeiss AxioImager Z1 motorized upright epifluorescence microscope (Carl Zeiss, Oberkochen, Germany), equipped with an Axiocam MR camera, and operated by the Zeiss Axiovision v4.7 software. Z-stacks (0.24/0.28 µm) covering the entire volume of individual mitotic cells were collected using a PlanApo 63×/1.40 NA or 40x/1.3 NA objective. Automated widefield microscopy was employed to assess nuclear circularity and micronuclei, and intensity levels of cGAS, γH2AX, Ki67, 53BP1/p21 and SA-β-gal biomarkers. Automated fluorescence microscopy was done using the IN Cell Analyzer 2000 (GE Healthcare), equipped with a Photometrics CoolSNAP K4 camera and using Nikon 20x/0.45 (immunostainings) or Nikon 40/0.95x (SA-β-gal assay) NA Plan Fluor objectives. Automated high-throughput image acquisition was also done using a Leica DMI6000 FFW inverted epifluorescence microscope (Leica Microsystems, Germany), equipped with a Hamamatsu FLASH 4.0 camera (Hamamatsu, Japan) and controlled by LAS X v2.0 software. Images were acquired using either a 40x/0.6NA dry objective or a 63x/1.30NA glycerol objective, both with appropriate filter cubes for each fluorophore. Identical settings were used for all matched images, and the minimum exposure time was used to avoid signal saturation. LaminA/C and LaminB1 immunostaining images were acquired using a Leica SP8 laser scanning confocal microscope (Leica Microsystems, Germany) with a 63x/1.30NA glycerol objective and controlled by the LAS X (v3.5.6 software, Leica SP8) software. DAPI (blue channel) and Lamin B1 (green channel) immunostainings were acquired using PMT detectors, whereas LaminA/C or γH2AX (red channel) staining was acquired using an HyD detector. Images were acquired with 0.33 µm z-stacks interval. Identical settings were used for all matched images, and the minimum laser power was used to avoid signal saturation. Histological images were acquired at different magnifications (skin, 4x; gonadal fat, 10x; aorta, 20x) using a Leica DM2000 LED microscope. Image deconvolution was done using AutoQuant X2 software (Media Cybernetics).

Phase-contrast long-term live-cell imaging

Fibroblasts grown in ibiTreat µ-Slides (ibidi GmbH) were imaged using a Leica DMI6000 epifluorescence microscope (Leica Microsystems, Germany) outfitted with an Orca Flash 4.0v2.0 camera (Hamamatsu, Japan), under conditions of controlled temperature, atmosphere, and humidity. Neighbor fields (30-40) were imaged every 2.5 min for 3 days, using a ×20 LD/0.4 NA objective (Leica Microsystems).

Fluorescence live-cell imaging

HGPS fibroblasts were cultured in ibiTreat 24-well µ-Slides (Ibidi GmbH) and labeled with SiR-DNA (251SC007, Spirochrome AG). Images were acquired with 10 min intervals under controlled environment using a Nikon Eclipse Ti motorized inverted epifluorescence microscope (Nikon), equipped with an Iris9 sCMOS camera (Photometrics) and a Spectra-X LED Light Engine (Lumencor), and using the 470/24 nm (10%) Spectra-X filter (Semrock) and a 40x/0.95 PL APO LAMBDA objective.

Image analysis

Live-cell and fixed-cell experiments were blindly quantified using the ImageJ/Fiji software. User-defined fluorescence intensity thresholds were set and consistently applied to samples within each experiment. α-tubulin intensity levels of calcium-stable k-fibers or astral microtubules were normalized to the mitotic spindle or cytoplasmic area, respectively, of each individual cell, and corrected for background. Interphase cells with DNA aggregates separated from the primary nucleus, but without apoptotic appearance, were considered to exhibit micronuclei. Threshold tool upon nucleus masking was used to measure nuclear circularity and background-corrected global fluorescence levels of LaminB1, Ku80, 5-FUrd, HeK9me3 and H4K20me3. For each independent experiment, values were normalized to the mean value of the control group. For nuclear morphology analysis, nuclei were considered normal if exhibiting smooth oval shape, and abnormal if exhibiting blebs, folds or gross irregularities in shape. For 53BP1+p21 and SA-β-gal staining, fluorescence intensity thresholds were established and applied uniformly to every sample in every experiment. In SA-β-gal activity assays, only cells displaying n > 5 fluorescent granules were considered positive. Cells positives for γH2AX double-strand breaks marker display two or more nuclear foci and S-phases were excluded. dsbNETs were analyzed using the 'Orthogonal views' function on ImageJ/FIJI, and quantified from 'Maximum Z-projection' images using ImageJ/FIJI. Cells were considered positive for dsbNETs if two or more tubules were observed extending deeper than the radius of the nucleus (*13*).

3D tracking of DNA DSBs

HeLa cells were seeded on glass bottom 8-well μ-Slides (ibidi GmbH) and transfected with pEGFP-h53BP1 plasmid two days before imaging. DSBs were induced the day before imaging, by treating cells overnight with 1 μM etoposide. Once cells expressing EGFP-53BP1-labelled foci were identified and imaging positions selected, the culture medium was replaced, and 100 nM UMK57, 100 nM UMK95 or 500 nM taxol was added. DSBs foci were imaged with a 40x/0.95 PL APO LAMBDA objective on a Nikon Eclipse Ti motorized inverted epifluorescence microscope (Nikon), equipped and an Iris9 sCMOS camera (Photometrics), a Spectra-X LED Light Engine (Lumencor) and a full-multiband filter set (Semrock). The 470/24 nm (10%) Spectra-X filter was used for imaging EGFP-53BP1. All imaging was done under controlled environment (37°C, 5% $CO^2$). Time-lapse recordings were done every 30 sec for 10 min, with 0.5 μm z-stacks intervals covering a range of 6 μm, using a binning of 2x2. Acquired images were preprocessed in Fiji/ImageJ by isolating each cell and subtracting background noise (radius 30px). Tracking analysis was performed in Imaris software (v9.6.1 Oxford Instruments) using the Imaris 3D Tracking Algorithm Autoregressive Model (0.5 μm max distance, no gaps). Spots identified objects with different sizes (starting diameter of 0.7 x 0.7 x 1.2 μm) through manual thresholding. An automatic filter was applied to remove small tracks, and poorly detected spots were removed by manual quality control. Tracking analysis was performed in a custom-made MATLAB (v. R2020) script, parsing the Imaris output and extracting each object's tracking positions over time. Initially, a registration step was conducted to compensate for live-cell movements: the centroid of all spots is subtracted for each time point from all the spots' positions. Subsequently, the tracks are organized into cell arrays by [time X Y Z] and normalized, so each track starts at a (0,0,0) coordinate. Finally, the mean-square displacement (MSD) was calculated using the msdanalyser (*44*) and the traveled distance of each spot was calculated using the Euclidean distance function.

RNA expression analysis (RT-qPCR)

Total RNA was extracted from both asynchronous and mitotic cell populations using Quick RNA MicroPrep (R1050, Zymo Research). Total RNA from snap-frozen samples of skin, fat, and aortic tissue was extracted by mechanical homogenization in a FastPrep-24 instrument (MP Biomedicals) using the RNeasy Fibrous Tissue Mini Kit (skin and fat) or the RNeasy Lipid Tissue Mini Kit (aorta) (both from QIAGEN) (fat and skin; QIAGEN). 0.8-1 μg of total RNA was reverse-transcribed using the NZY First-Strand cDNA Synthesis Kit (NZYTech). RT-qPCR was performed using the iTaq™ Universal SYBR® Green Supermix in a CFX96/384 Touch™ Real-

Time PCR Detection System (Bio-Rad Laboratories). Relative expression levels were determined using the CFX Maestro Software (Bio-Rad Laboratories). Primers used are listed in Tables S5 and S6.

Targeted transcriptome sequencing

10 ng of RNA was reverse-transcribed using the AmpliSeq Whole Transcriptome primers supplied in SuperScript VILO cDNA Synthesis kit (Thermo Fisher Scientific). The resulting cDNA was then used for targeted amplification (12 cycles) using Ion AmpliSeq primers and technology. Barcoded adapters were added and ligated to individual reactions according to the Ion AmpliSeq™ Transcriptome Human Gene Expression Kit (Thermo Fisher Scientific) instructions. The pooled libraries were processed on the Ion Chef™ System, and the resulting 550™ chip was sequenced on the Ion S5™ XL System (both from Thermo Fisher Scientific). Data were processed using the Ion Torrent platform-specific pipeline software, Torrent Suite v5.8.0, which generated sequence reads, trimmed adapter sequences, filtered and removed poor signal reads, and split the reads according to the barcode. The FASTQ and/or BAM files generated by the Torrent Suit plug-in FileExporter v5.12 were analyzed using Torrent Suite™ v5.8.0 Software (Thermo Fisher Scientific) running Ion AmpliSeq™ RNA plug-in v5.12, coverage Analysis plug-in v5.12, and target region hg19_AmpliSeq_Transcriptome_21K_v1. Differential gene expression analysis was performed using the Transcriptome Analysis Console v4.0.2 (TAC) Software (Thermo Fisher Scientific). Gene expression was defined as significantly different based on $p < 0.05$ for the comparison between UMK57 versus DMSO-treated DS and HGPS fibroblasts. The targeted transcriptome sequencing data represents two independent experimental replicates of two biological samples from HGPS and DS.

Functional Enrichment Analysis

For the identification of processes and signaling pathways modulated by UMK57 treatment, GSEA was performed for transcriptomic data based on a pre-ranked list of differentially expressed genes for each condition (HGPS and DS). The metric of each differentially expressed gene was "1" for upregulated genes and "-1" for downregulated genes upon UMK57 treatment. The enrichment analysis was executed in R (v4.2.3) with the GSEA function from the cluster Profiler package (v4.6.2) together with the fgsea package (v1.24.0) using the hallmark and gene ontology biological processes gene sets obtained from MSigDB using the msigdbr package (v7.5.1, maximum gene set size = 1000). *p*-values were corrected for multiple comparisons using the Benjamini-Hochberg method, and a *p* adjust cutoff of 0.1 was used to select statistically significant terms. For the identification of enriched biological processes in common upregulated genes from HGPS and DS fibroblasts following UMK57 treatment, the enrichGO function from clusterProfiler was used. Statistically significant processes (*p* adjust < 0.1) were obtained after false-discovery rate correction, and the treeplot function was used for hierarchical clustering of relevant terms.

Western blotting

Protein samples were collected from cell pellets extracted in RIPA lysis buffer with protease inhibitors (89900, ThermoFisher Scientific). Protein content was determined by the Lowry method (DC™ Protein Assay, Bio-Rad Laboratories) following the manufacturer's instructions. Equal amount of protein extracts were loaded for SDS-polyacrylamide gel electrophoresis and transferred to nitrocellulose membranes for Western blot analysis. Blocking was performed with 5% non-fat dry milk in TBS-T (50 mM Tris-HCl pH=7.4, 150 mM NaCl, 0.05% Tween-20) for 1 h. Membranes were then incubated overnight at 4ºC with the indicated antibodies. Both, primary and secondary antibodies were diluted in TBS-T + 2% non-fat milk as follows: rabbit anti-KIF2C

(#sc-81305, Santa Cruz,1:500), rabbit anti-γH2AX (#9718, Cell Signaling Technology, 1:1,000), rabbit anti-cGAS (15102, Cell Signaling Technology, 1:1,000), mouse anti-GAPDH (#60004, ProteinTech Group, 1:30,000), mouse anti-α-tubulin (T5168, Sigma-Aldrich, 1:100,000), HRP-conjugated goat anti-rabbit (111-035-003, Jackson Immunoresearch, 1:8,000) and HRP-conjugated goat anti-mouse (115-035-003, Jackson Immunoresearch, 1:8,000). HRP conjugates were detected using Clarity Western ECL Substrate reagent (Bio-Rad Laboratories) according to the manufacturer's instructions. A GS-900 calibrated densitometer operated by the ImageLab™ software v6.0.1 (Bio-Rad Laboratories) was used for quantitative analysis of protein levels.

Mouse strains and compliance with ethical principles

Procedures involving animals and their care were conducted with institutional ethical guidelines (i3S Animal Welfare and Ethics Review Body, ORBEA) and with the National and European Union regulations (DL 113/2013 and 2010/63/EU, respectively), under DGAV license (DGAV 0421/000/000/2017). Mice were maintained under a 12 h light/dark cycle and fed with regular sterilized rodent's chow (Envigo 2014S) and tap water *ad libitum*, at the i3S animal facility. Mice carrying the Lmna mutation p.Gly609Gly ($Lmna^{G6096G/G609G}$ in C57BL/6 background) referred to as LAKI (*28*) were used as study model. Experiments were performed using littermates of both sexes, randomly assigned to control (DMSO) and experimental (UMK57) therapy. Regarding the control group, we selected $Lmna^{+/+}$ homozygous mice (referred to as wild-type) from the same genetic background as LAKI. Extended information on mouse housing and husbandry, can be found in the Supplementary Notes section.

UMK57 treatment *in vivo*

UMK57 was administered subcutaneously, using osmotic pumps (ALZET® Osmotic Pumps, model #1004), which allow controlled drug delivery at a rate of 0.11 µL/hr for 4 weeks. UMK57 was dissolved in a 50% DMSO / 50% PEG400 solution at a concentration of 17.5 mM. UMK57 was administered for 8 weeks (0.7 mg/kg/day) via sequential implant of two osmotic pumps. The first pump was implanted in 5-week-old animals, removed after 4 weeks, and replaced by a new pump. Animals were euthanized 4 weeks later (13-week-old). This route of administration circumvented the poor solubility (Table S1), moderate permeability (Table S2) and fast clearance (Table S3) constrains of the UMK57 drug as determined by assays outsourced to Cyprotex Discovery Ltd (Macclesfield, UK).  UMK57 biodistribution was confirmed by liquid-chromatography mass-spectrometry (LC-MS) analysis of several tissues collected at the end of the treatment (outsourced to Cyprotex Discovery Ltd).

Grip-strength test

Grip strength was measured using a BIO-GS3 (Bioseb). Mice were placed on the grating with the two front paws attached and gently pulled back to measure grip strength until releasing from the grid. The values obtained represent the muscle force (g) obtained in n=2 trials per animal with an interval of 5 min.

Cytokine array

Cytokine levels were measured in mouse serum sample replicates using the Mouse High Sensitivity T Cell 18-Plex Discovery Assay Array (MDHSTC18; Eve Technologies).

Histological analysis

Tissue biopsies of skin, gonadal fat, and aorta were collected, fixed overnight in 10% (v/v) buffered formalin, and routinely processed in an automated tissue processor for paraffin embedding. Serial 4µm-thick sections were prepared and stained with hematoxylin and eosin (H&E) using standard procedures.

Three-dimensional X-ray microcomputed tomography

X-ray microcomputed tomography was performed using SkyScan 1276 (Bruker). The scanning parameters were as follows: image pixel size of 40/10 µm, 70 kV, 200 µA, 0.5 mm aluminum filter, image averaged over 2 frames, 180° rotation, and 0.8° rotation step. 3D reconstructions of tomographic slice images were processed and analyzed using Bruker software (NRecon v1.7.5.0, DataViewer v1.5.6.3, CTAn v1.20.3.0, CTVox v3.3.0-1412, and CTVol v2.3.2.1). Quantification of the kyphosis index was as described in (*45*). Total fat volume was quantified from a standardized and referenced set of cross-sectional slices. For bone microarchitecture analysis, cross-sectional slices of trabecular (metaphyseal) tibial bones were selected concerning the growth plate, following the American Society for Bone and Mineral Research (ASBMR) parameters. This analysis assessed bone volume fraction (BV/TV), the number of trabeculae per unit length (Tb.N), trabeculae separation (Tb.Sp), trabeculae thickness (Tb.Th), and bone mineral density (BMD).

Statistical analysis

All data presented are at least from two independent experiments and are shown as mean ± s.d or median ± range. Statistical analyses were performed using Prism® 8 software (GraphPad). Sample sizes are indicated in each graph. Normal distribution of the data was tested using the Shapiro–Wilk test, and the appropriate statistical test was used according to the data distribution and mentioned in the respective figure legend. All relevant *p*-values are shown in the figures, where $p<0.05$ is considered as significant.

### Supplementary Text

Mouse housing and husbandry

Mice were maintained in a specific pathogen-free (SPF), AAALAC-accredited barrier facility at i3S – Instituto de Investigacão e Inovacão em Saúde, on a 12h light:12h dark cycle, at 20-24ºC, 45-55% humidity and positive air pressure. i3S animal facility performs regular monitoring of known viral, bacterial, and parasitic pathogens. All animals had specific pathogen-free sanitary status (SPF), according to the FELASA recommendations. Mice were housed at a preferred stocking density of 3-4 mice per cage, in Eurostandard Type II cages (1264C, Tecniplast - 268 x 215 x 141 mm, floor area 370 $cm^2$) with a filter top-lid. In addition to corn cob bedding substrate (changed weekly; Scobis Due, Mucedola), standard environmental enrichment was provided, consisting of an E-Cube (Allentown), one clear handling tube (Datesand), one mouse cellulose shelter (LBS), and wooden chew blocks (Aspen bricks, Sodispan Research). Pups were PCR-genotyped at day 8, separated by sex, and littermates were housed together and established into experimental cohorts according to the single mutant mouse strain ($Lmna^{G6096G/G609G}$ in C57BL/6 background). Mice were given sterilized water and diet ad libitum (A155D00623 - Complete feed with low phytoestrogen content/soybean free - 10 kJ% fat, 23 kJ% protein, 67% kJ carbohydrates). Gel recovery and moistened supplemental food were added to the bottom of each cage after surgery to facilitate food consumption.

Extended experimental procedures on live animals

In this study, both females and males were weighed regularly (bi-weekly monitoring). Animals were humanely euthanized if they showed ≥20% bodyweight loss and/or the presence of clinical, behavioral, and physiological signs (as defined in Directive 2010/63/EU). Animals used in experiments were not allowed to breed.

For 3D micro-CT *in vivo* analysis (body fat scanning), mice were anesthetized in an induction chamber, using 4-5% isoflurane with an oxygen source, and were then transferred to the SkyScan 1276 device (Bruker, Belgium) (temperature: 37ºC). Ophthalmic ointment was applied to both eyes to prevent dryness and corneal damage. Anesthesia (1-2% isoflurane) was maintained throughout scanning. Animals were allowed to recover alone in a clean cage and then returned to the home cage when fully recovered.

To measure grip-strength, mice were placed on the grid with all two paws attached and gently pulled backward. Each animal was subjected to 2 measurements within 5 minutes of resting in the home cage.

To obtain tissue samples, mice were anesthetized with isoflurane (volatile anesthetic) prior to death by decapitation. Dissection was performed as soon as possible after euthanasia, with the organs being removed, sectioned, and either snap-frozen in liquid nitrogen (stored at -80ºC) or placed in formalin for fixation (at room temperature).

Blood collection was performed by retro-orbital under volatile isoflurane anesthesia, followed by euthanasia by decapitation.

### Supplementary Figures

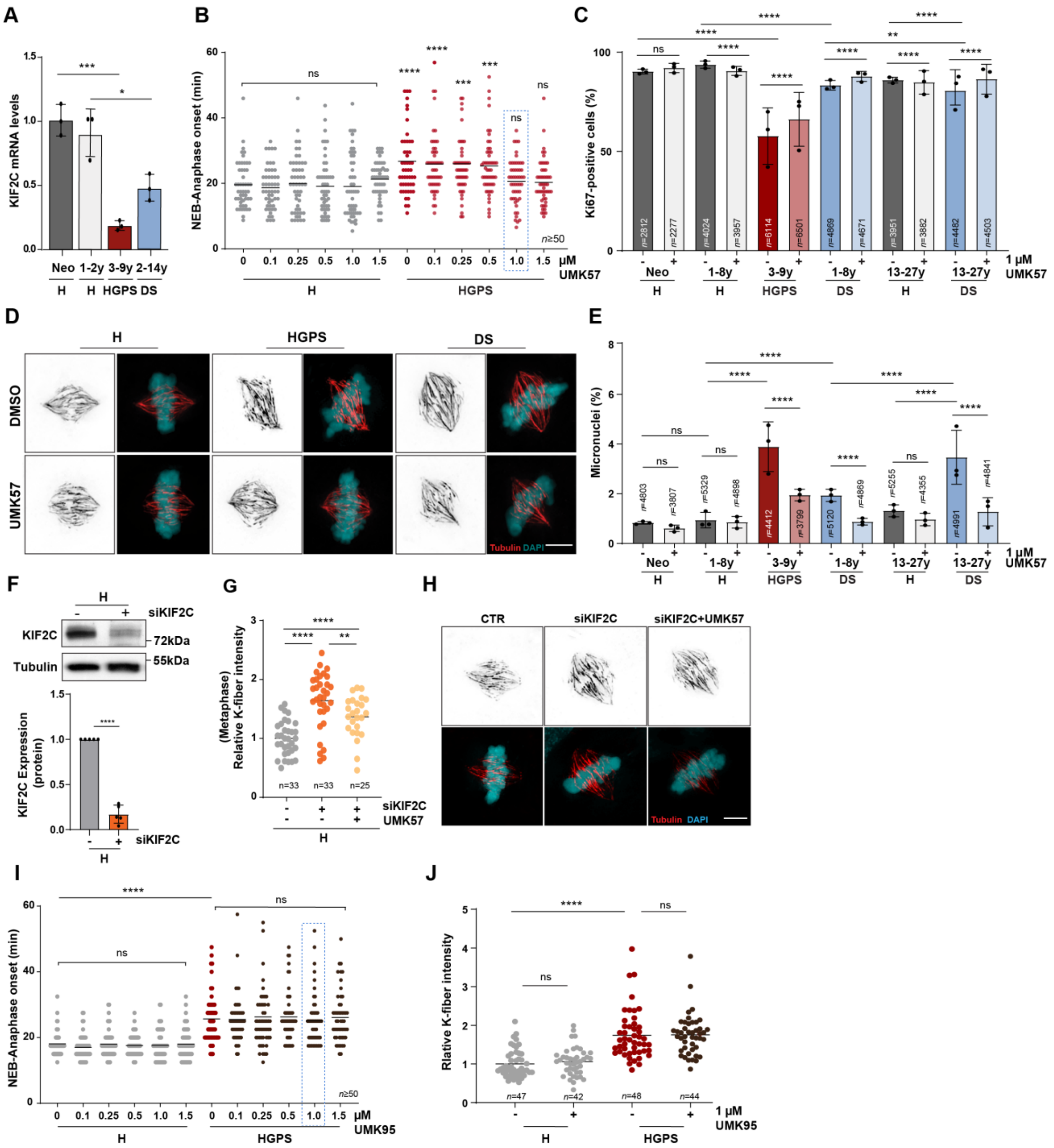


**Fig. S1. Titration of UMK57 optimal dose and validation of its specific effect on the restoration of k-MT dynamics via KIF2C targeting (related to Fig. 1).**

**A)** *KIF2C* transcript levels ($2^{-\Delta\Delta Ct}$) in mitotic extracts from healthy (H), HGPS, and DS fibroblasts, normalized to *HPRT1* and *GAPDH* housekeeping genes, and using neonatal control as reference.

**B)** H and HGPS fibroblasts were treated for 24 hours with increasing concentrations of UMK57 to determine the optimal dose capable of rescuing the mitotic delay (due to defective k-MT dynamics) in HGPS cells, while having no effect on the mitotic duration (nuclear envelope breakdown to anaphase onset) of healthy controls. UMK57 was used at 1 μM for all subsequent experiments. **C)** Percentage of Ki67-positive cells in H, HGPS, and DS cultures treated with DMSO or UMK57. **D)** Representative images and quantification of calcium-stable k-fiber intensity by immunofluorescence analysis of tubulin-stained (red) metaphase cells treated with DMSO (-) and UMK57 (+) as indicated. DNA is shown in cyan. Levels were compared to DMSO-treated neonatal cells. **E)** Percentage of H, HGPS, and DS cells with micronuclei upon treatment with DMSO (-) or UMK57 (+). **F)** Western blot analysis of KIF2C protein levels upon siRNA-mediated knockdown of KIF2C (siKIF2C) in H cells. Tubulin was used as loading control, and protein levels were normalized to mock control (-). **G, H)** Quantification (G) and representative images (H) of calcium-stable k-fiber intensity levels in mock or siKIF2C-depleted metaphase cells treated with UMK57 or DMSO. DNA is shown in cyan. Levels were normalized to untreated control. **I)** Unchanged mitotic duration in H and HGPS fibroblasts treated with increasing concentrations of an inactive drug analog (UMK95) for 24 h. UMK95 was used at 1 μM for all subsequent experiments. **J)** Quantification of calcium-stable k-fiber intensity levels in H and HGPS metaphase cells treated with DMSO (-) or UMK95 (+). Levels were normalized to untreated H control. Scale bars, 5 μm. All values are the mean ± s.d. of at least three independent experiments. The sample size (n) is indicated in each graph. ns $p>0.05$, * $p<0.05$, ** $p<0.01$, *** $p<0.001$, **** $p<0.0001$ by Mann-Whitney (B, G, I, J), Fisher's exact (C, E) and Unpaired t-test (A, F) statistical tests.

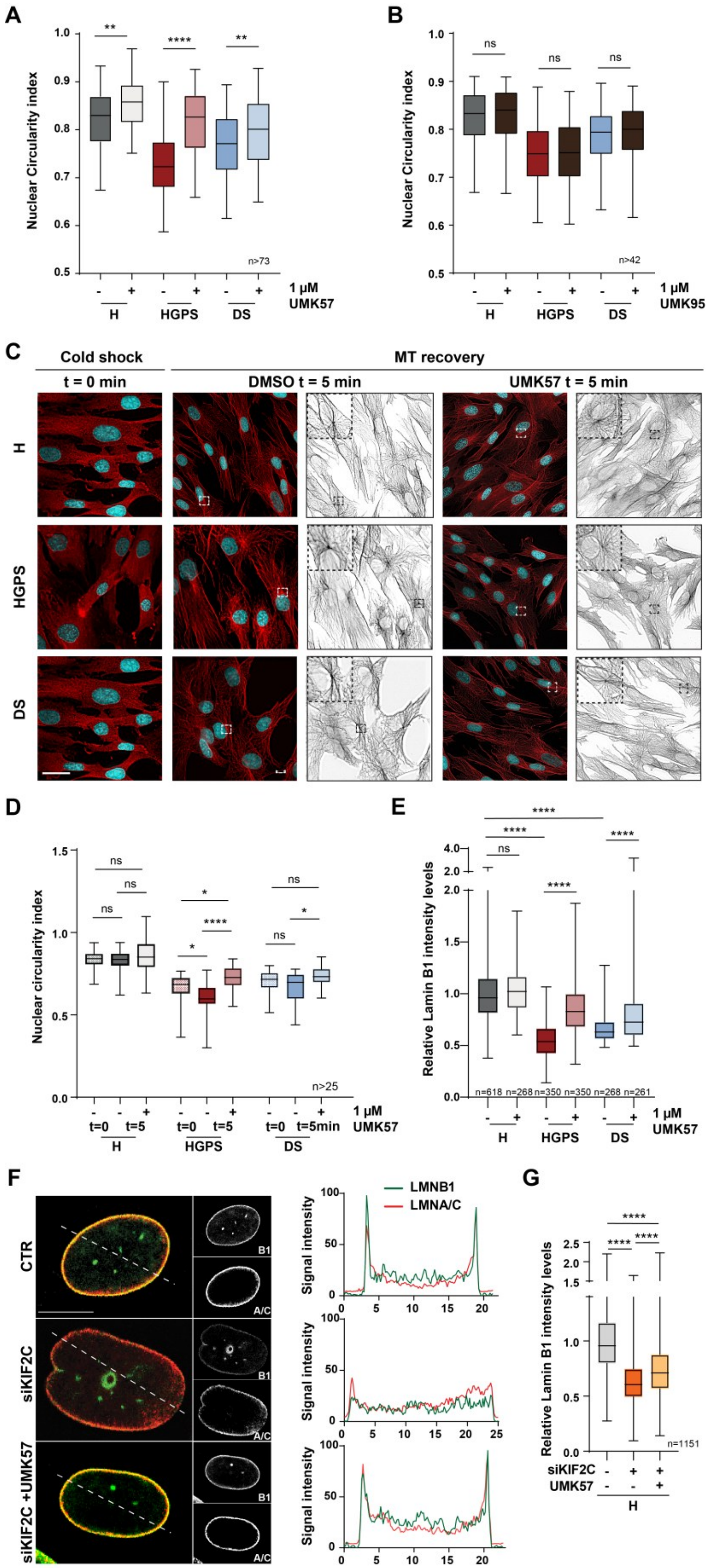

A
Nuclear Circularity index
n>73
1 µM UMK57
H
HGPS
DS
B
n>42
1 µM UMK95
C
Cold shock
t = 0 min
MT recovery
DMSO t = 5 min
UMK57 t = 5 min
D
Nuclear circularity index
n>25
E
Relative Lamin B1 intensity levels
n=618 n=268 n=350 n=350 n=268 n=261
F
CTR
siKIF2C
siKIF2C +UMK57
B1
A/C
LMNB1
LMNA/C
Signal intensity
G
n=1151
siKIF2C
UMK57

**Fig. S2. Nuclear dysmorphology and laminopathy in HGPS and DS cells are rescued by specific targeting of KIF2C motor activity by the UMK57 drug agonist (related to Fig. 1).** **A, B)** Quantification of nuclear circularity index in H, HGPS and DS cells upon UMK57 treatment (A) and UMK95 treatment (B). **C, D)** Representative immunofluorescence images (C) and nuclear circularity index (D) measured in H, HGPS and DS cells upon microtubule (MT) depolymerization induced by cold treatment (t=0 min) and MT regrowth (t=5 min) in the absence (-) or presence (+) of UMK57. **E)** Quantification of LMNB1 fluorescence intensity levels in H, HGPS and DS cells treated with DMSO (-) or UMK57 (+). **F)** Representative images and histogram profiling of LMNA/C (red) and LMNB1 (green) nuclear intensity levels in mock and siKIF2C-depleted cells treated with DMSO or UMK57. **G)** Quantification of LMNB1 fluorescence intensity levels in mock and siKIF2C-depleted cells treated with DMSO or UMK57. Scale bar, 5 μm (C) and 10 μm (F). All values are the median ± range of at least three independent experiments. The sample size (n) is indicated in each graph. ns $p>0.05$, * $p<0.05$, ** $p<0.01$, *** $p<0.001$, **** $p<0.0001$ by Mann-Whitney (A, B) and Kruskal-Wallis test followed by Dunn's multiple comparisons test (D, E, G) statistical tests.

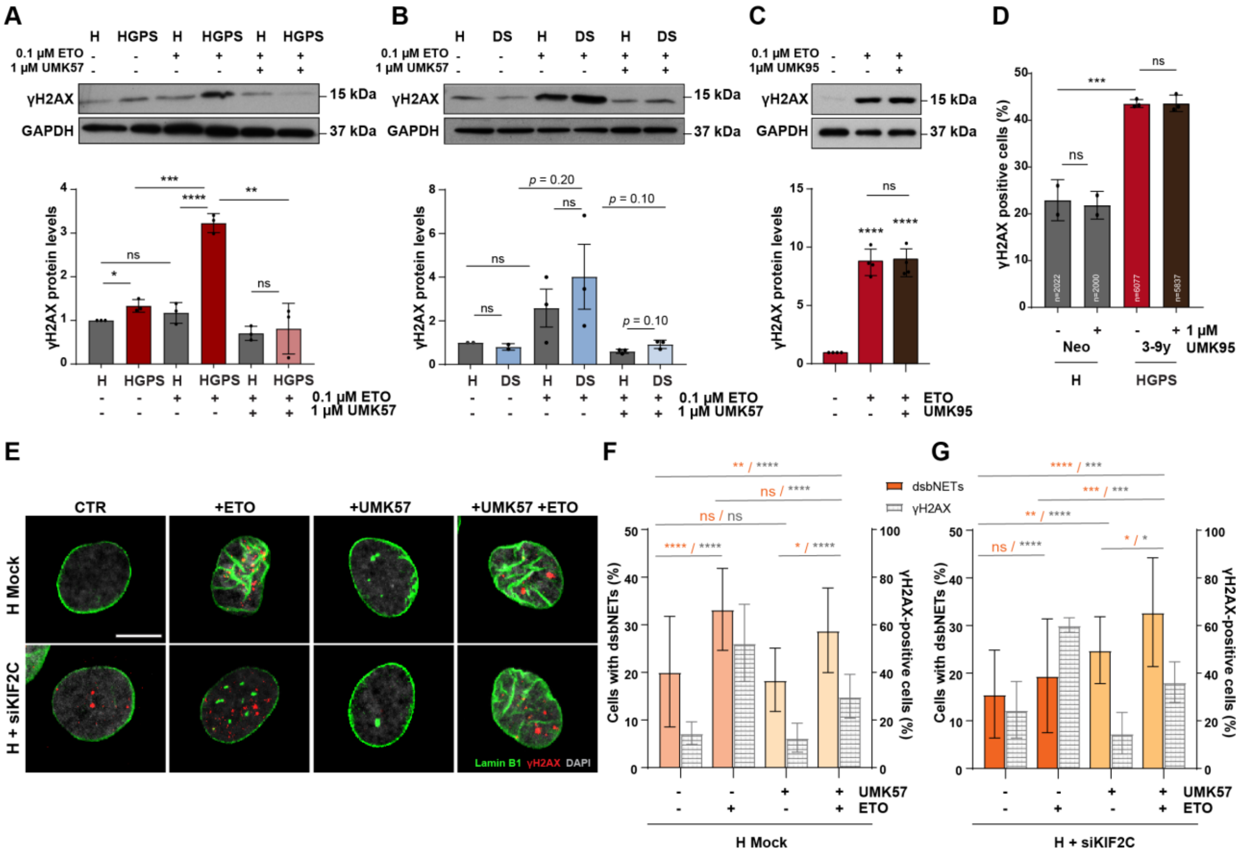


**Fig. S3. Pharmacological enhancement of KIF2C activity specifically boosts DSB DNA repair capacity (related to Fig. 2).**

**A, B)** Western blot analysis of γH2AX protein levels in total extracts from HGPS and DS fibroblasts, untreated or challenged with the DSB-inducing agent etoposide, in the absence or presence of UMK57. GAPDH was used as a loading control. **C, D)** Western blot (C) and immunofluorescence (D) analysis of γH2AX protein levels/positive cells in healthy controls treated with etoposide and UMK95 as indicated. GAPDH was used as loading control. **E)** Representative images of LaminB1/dsbNETs (green) and γH2AX/DSBs (red) immunostainings used for quantitative analysis. **F, G)** Quantification of dsbNETs-positive cells (left axis) and γH2AX-positive cells (right axis) in mock (F) and siKIF2C-depleted (G) H cells challenged with etoposide (ETO, 10 µM) after treatment with UMK57 or DMSO (n > 250 cells). Scale bar, 5 µm. All values are the mean ± s.d. of at least two independent experiments. The sample size (n) is indicated in each graph. ns $p>0.05$, * $p<0.05$, ** $p<0.01$, *** $p<0.001$, **** $p<0.0001$ by unpaired t-test (A), Mann-Whitney (B), Fisher's exact test (F, G) and ordinary one-way ANOVA with Tukey's multiple comparison correction (C, D) statistical tests.

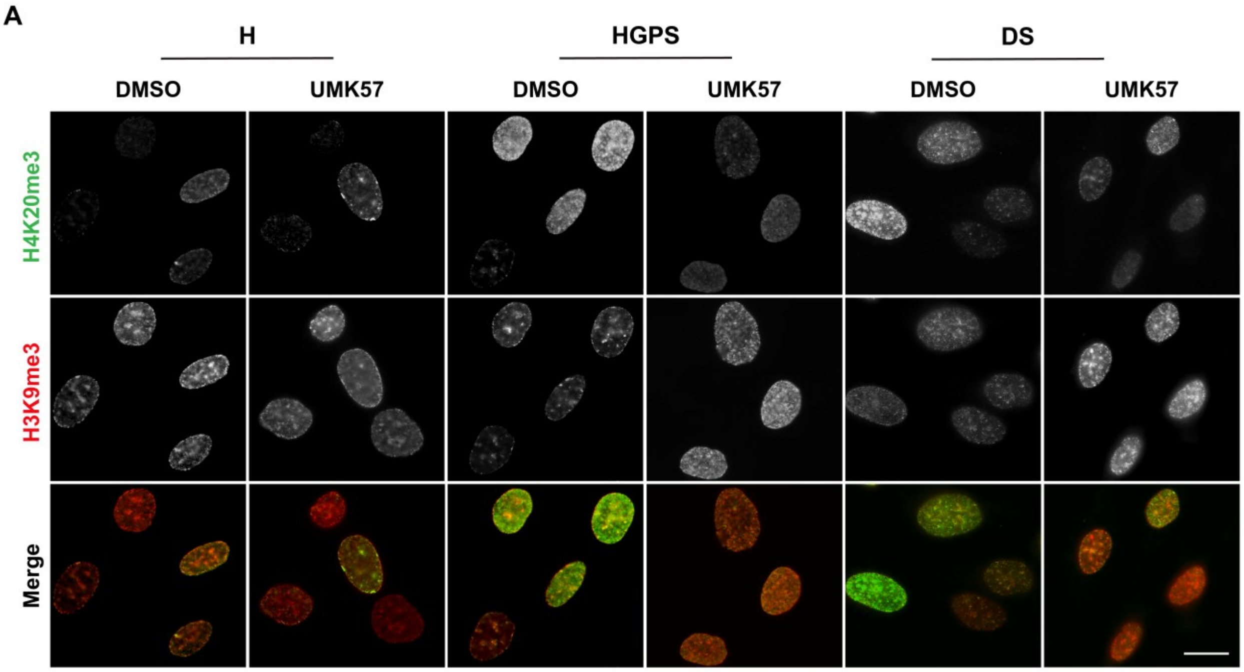


**Fig. S4. Restored heterochromatin-associated histone epigenetic marks upon treatment of HGPS and DS cells with UMK57 (related to Fig. 3).**

**A)** Immunofluorescence of heterochromatin-associated epigenetic marks H3K9me3 (red) and H4K20me3 (green) in fibroblasts from healthy (H), Down Syndrome (DS) and Hutchinson-Gilford progeria syndrome (HGPS) donors, treated with DMSO and UMK57. Scale bar, 20 µm.

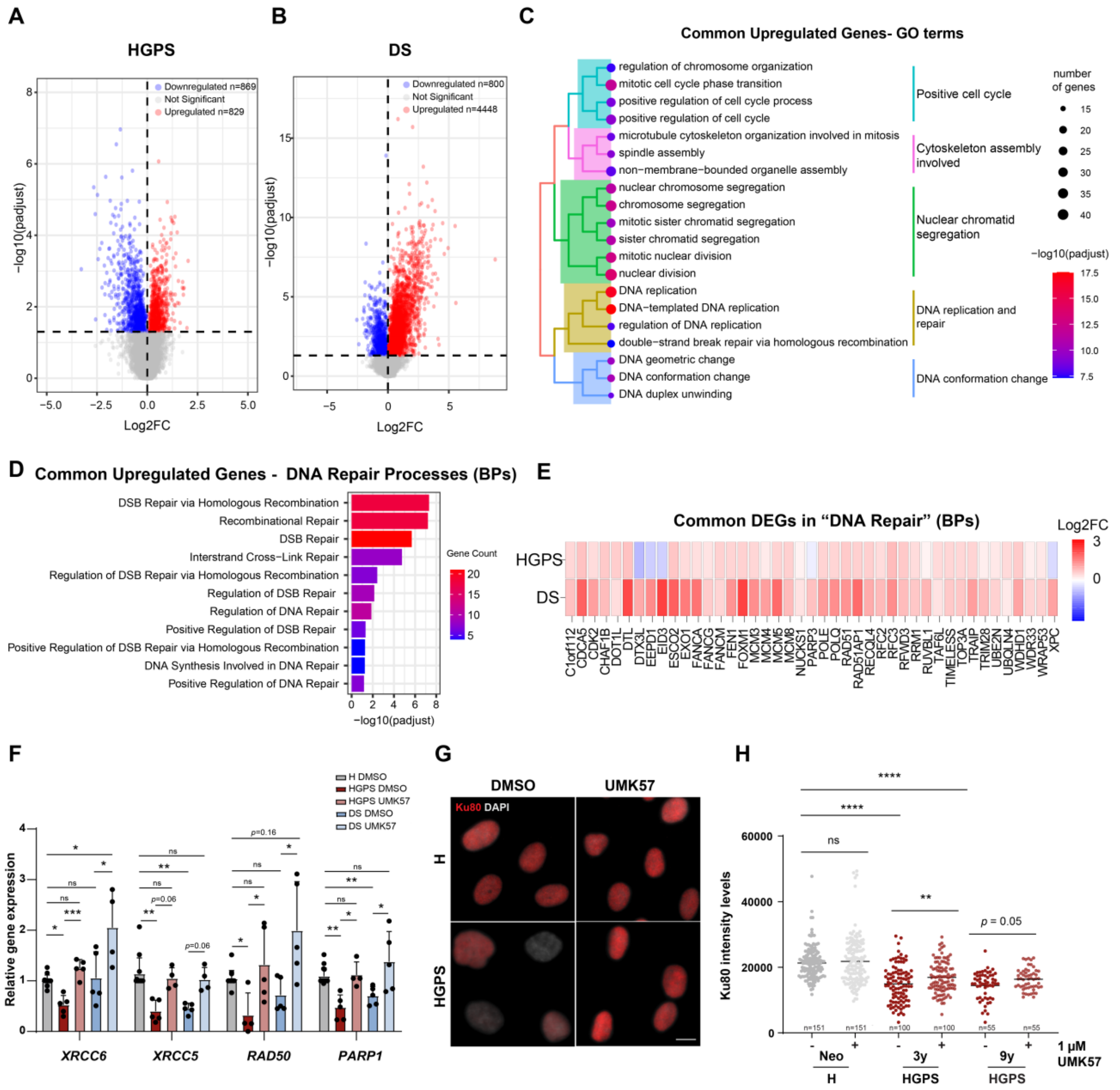


**Fig. S5. KIF2C activity restores favorable chromatin organization, thereby enhancing DNA repair pathways in cells affected by HGPS and DS (related to Fig. 3).**

**A, B)** Volcano plots illustrating changes in individual gene expression following UMK57 treatment of HGPS (A) and DS (B) fibroblasts. Significantly upregulated and downregulated genes in red and blue, respectively. **C)** Tree plot showing hierarchical clustering of enriched GO terms for upregulated genes found in both HGPS and DS cells treated with UMK57 (biological processes). **D)** Gene ontology analysis of DNA repair pathways (from MSigDB C5 - biological processes) that are significantly enriched in both HGPS and DS cells following UMK57 treatment. BPs are plotted accordingly to descending order of significance. **E)** Heatmap of DNA repair genes (GO:0006281) significantly upregulated by UMK57 treatment in both HGPS and DS fibroblasts. Color intensities represent log2FC. **F)** Transcript levels of DNA repair genes in healthy (H) and

HGPS or DS fibroblasts treated with DMSO or UMK57 for 48 h. *TBP* and *HPRT1* were used as reference genes. All levels were normalized to the DMSO-treated healthy sample. **G)** Representative images of H and HGPS fibroblasts treated with DMSO or UMK57 and immunostained for XCRCC5/KU80. **H)** Quantification of KU80 intensity levels by immunofluorescence analysis. Scale bar, 10 µm. All values are the mean ± s.d. of at least three independent experiments. The sample size (n) is indicated in each graph. ns $p$>0.05, * $p$<0.05, ** $p$<0.01, *** $p$<0.001, **** $p$<0.0001 by Kruskal-Wallis test followed by Dunn's multiple comparisons test (F) and Mann-Whitney (H) statistical tests.

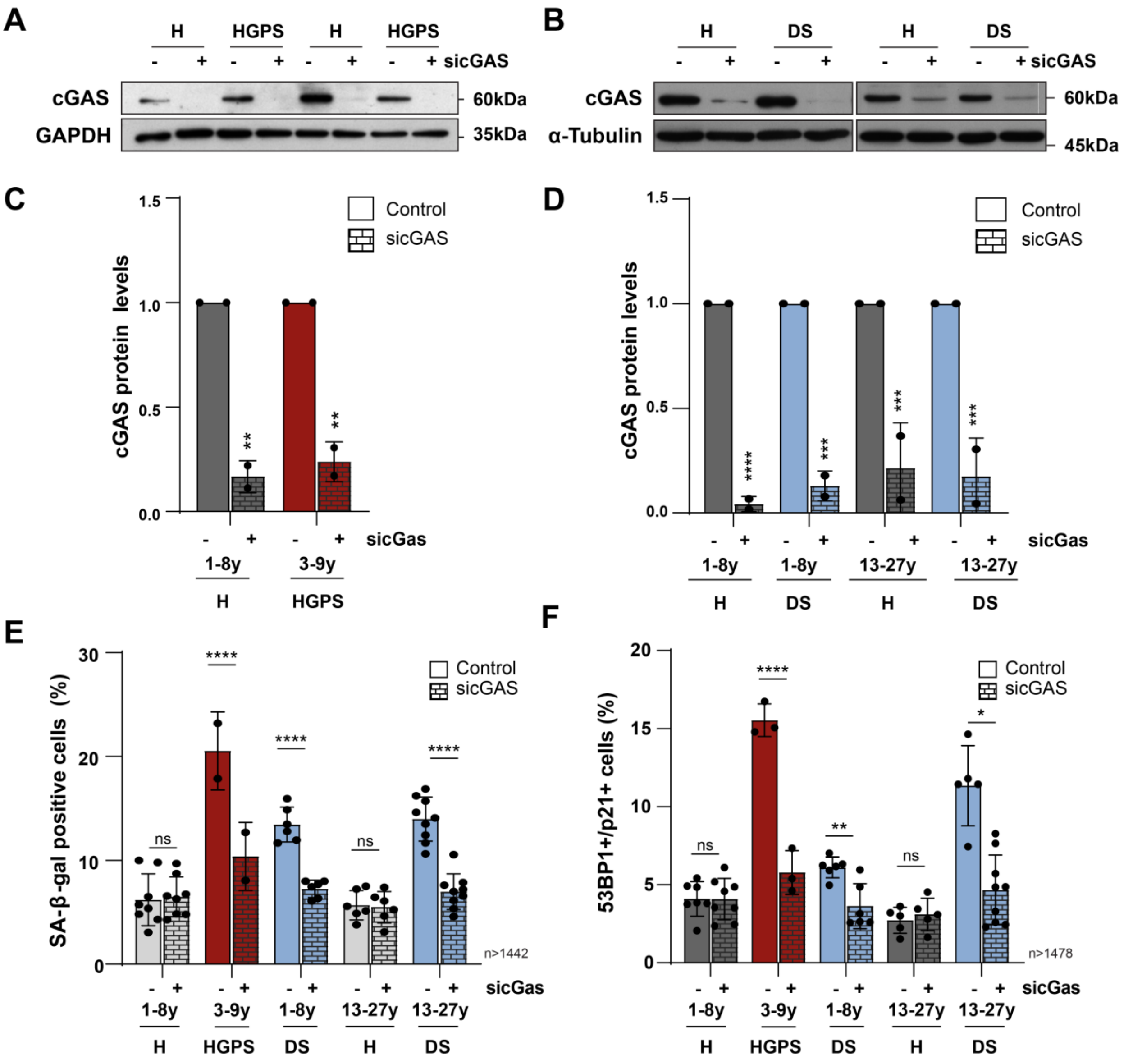


**Fig. S6. cGAS-STING signaling pathway is a key mediator of senescence accrual in HGPS and DS cell cultures (related to Fig. 3).**

**A-D)** cGAS protein levels following siRNA-mediated depletion (sicGAS) in fibroblast cultures from healthy (H), HGPS (A,C) and DS (B,D) donors. Protein levels were normalized to the loading controls (GAPDH or tubulin). Untreated H controls were used as reference. **E)** Percentage of cells with SA-β-galactosidase activity and **F)** staining double-positive for Cdkn1a/p21 cell cycle inhibitor and 53BP1 DNA damage (≥1 foci) senescence markers after siRNA-mediated depletion of cGAS. All values shown are mean ± s.d. of at least two independent experiments. The sample size (n) is indicated in each graph. ns $p$>0.05, * $p$<0.05, ** $p$<0.01, *** $p$<0.001, **** $p$<0.0001 by unpaired t-test (C, D) and Fisher's exact test (E, F) statistical tests.

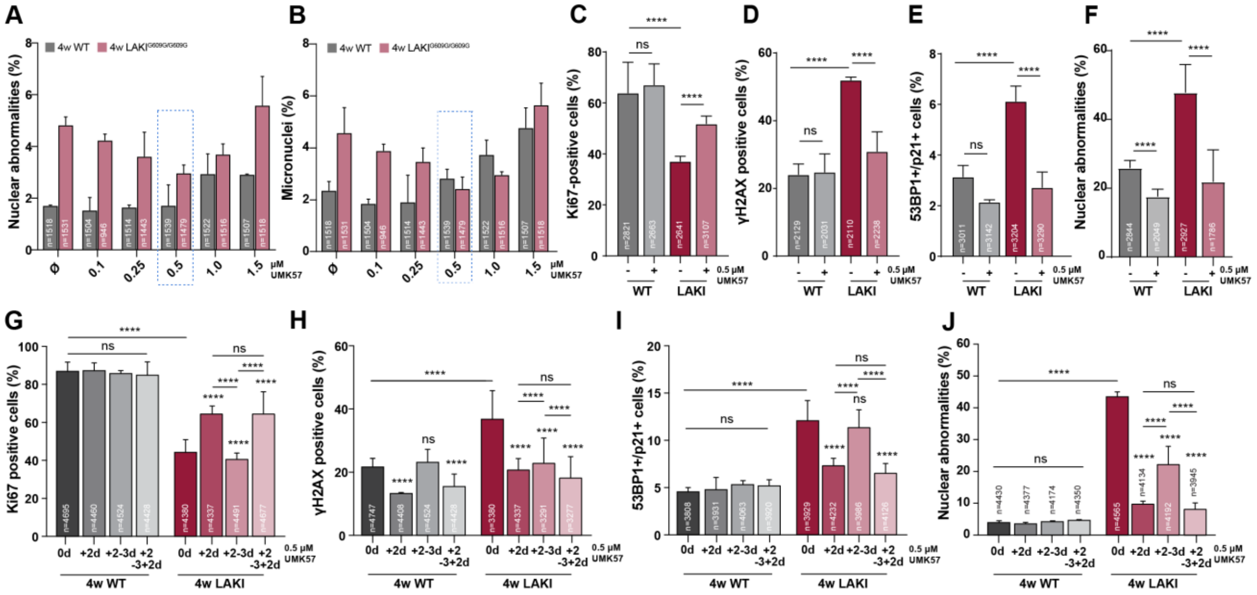


**Fig. S7. Evaluation of senescence features mitigated by UMK57 treatment in LAKI mouse fibroblasts (related to Fig. 4).**

**A, B)** Titration of UMK57 optimal dose by evaluation of rescue efficacy of nuclear abnormalities (A) and micronuclei (B) in murine adult fibroblasts (MAFs) from 4-week-old LAKI mice upon 4 days of treatment (MAFs from 4-week-old WT mice used as controls). UMK57 was used at 0.5 uM in all subsequent experiments. **C-F)** Percentage of WT or LAKI MAFs staining positive for the Ki67 proliferation marker (C), γH2AX DNA damage marker (D), Cdkn1a/p21 cell cycle inhibitor and 53BP1 DNA damage senescence markers (E), and exhibiting nuclear abnormalities (F) upon treatment with UMK57 or DMSO for 4 days. **G-J)** Cyclic drug treatment of WT and LAKI MAFs consisting in 2 days treatment with UMK57, followed by 3 days of drug withdrawal, and again 2 days treatment with UMK57. Evaluation of treatment efficacy by quantitative analysis of the percentage of cells staining positive for Ki67 (G), γH2AX (H) and Cdkn1a/p21 + 53BP1 (I), and exhibiting nuclear abnormalities (J). All values shown are mean ± s.d. of at least two independent experiments. The sample size (n) is indicated in each graph. ns $p>0.05$, * $p<0.05$, ** $p<0.01$, *** $p<0.001$, **** $p<0.0001$ by and Fisher's exact test statistical tests.

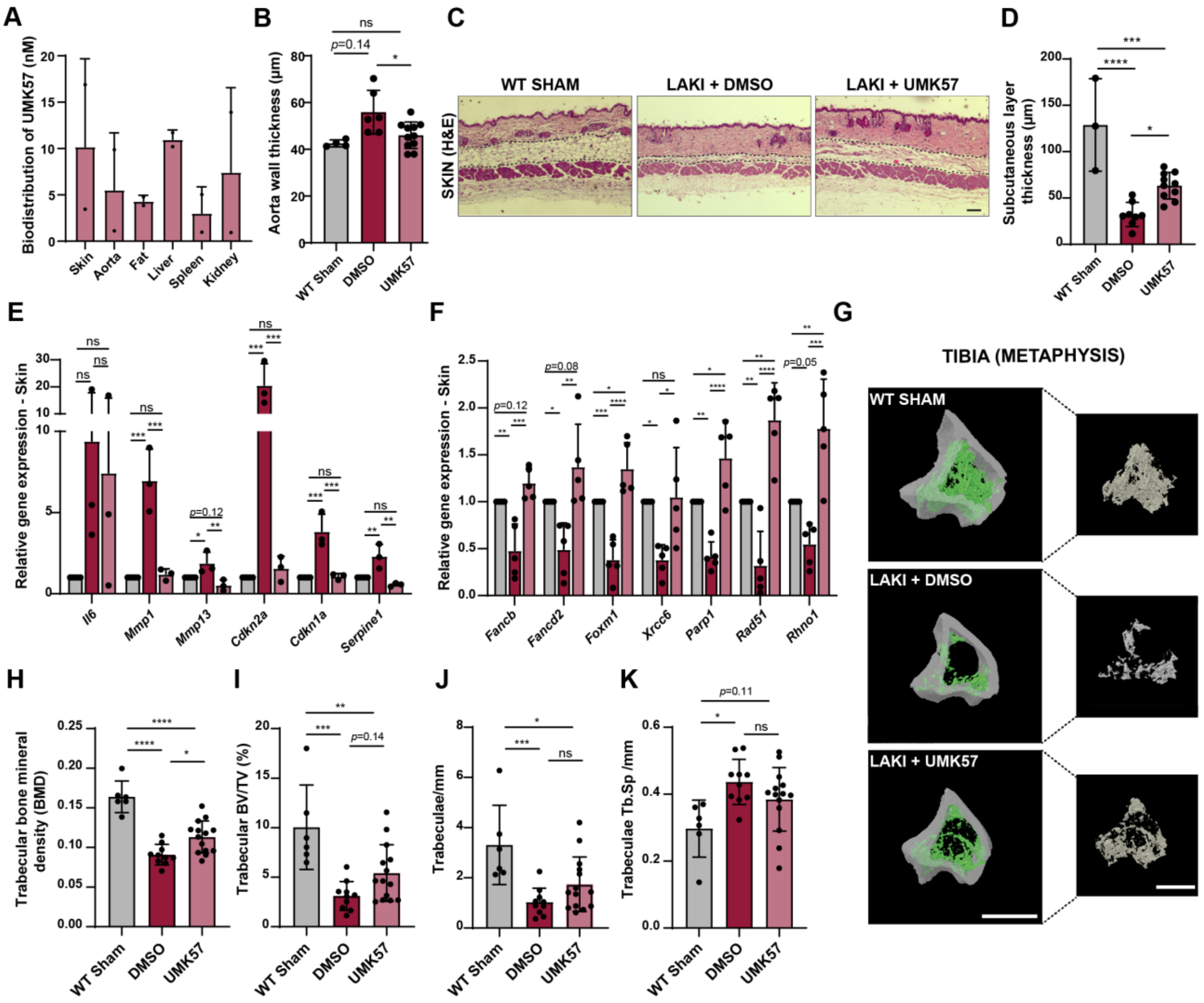


**Fig. S8. Improvement of histopathological features and associated molecular signatures upon UMK57 treatment of LAKI mice (related to Fig. 4).**

**A)** LC-MS/MS analysis of UMK57 biodistribution across tissues collected from LAKI mice treated via osmotic pumps. **B)** Aortic wall thickness measured in histological sections from WT mice and LAKI mice treated with DMSO and UMK57 as shown in Fig. 4M. **C)** Histological analysis of telogenic skin in WT mice and in LAKI mice treated with DMSO or UMK57. Dashed lines delineate the hypodermal adipose layer. **D)** Hypodermis layer thickness in telogenic skin sections as shown in C. **E)** RT–qPCR analysis of senescence gene expression levels ($2^{-\Delta\Delta Ct}$) in the skin. **F)** RT–qPCR analysis of DNA repair gene expression levels ($2^{-\Delta\Delta Ct}$) in the skin. **G)** micro-CT analysis of the trabecular tibia (metaphysis). **H)** Quantification of trabecular bone mineral density (BMD), **I)** bone volume fraction (BV/TV), **J)** number of trabeculae (Tb.N), **K)** and trabeculae thickness (Tb.Sp) from WT and LAKI mice treated with DMSO or UMK57. Scale bar, 1 mm. All values are the mean ± s.d. of at least three independent experiments. The sample size (n) is indicated in each graph. ns $p>0.05$, * $p<0.05$, ** $p<0.01$, *** $p<0.001$, **** $p<0.0001$ by Kruskal-Wallis test followed by Dunn's multiple comparison test (B, D, H-K) and ordinary one-way ANOVA parametric test with Tukey's multiple-comparison correction (E, F) statistical tests.

**Table S1.** Thermodynamic solubility of the UMK57 compound (provided by Cyprotex Discovery, Macclesfield, UK).

| **Compound** | **Thermodynamic solubility** | | | | | |
|---|---|---|---|---|---|---|
| **Name** | **Solubility (mg/mL)** | | **Mean solubility** | **SD** | **n** | **Variables** |
| | Replicate 1 | Replicate 2 | | | | |
| metoprolol | >5.00 | >5.00 | NC | NC | 2 | Temperature=RoomTemp IncubationTime=Overnigh DetectionMethod=HPLC/UV |
| nicardipine | NC | NC | NC | | | |
| pyrene | NC | NC | NC | | | |
| **UMK57** | <0.0250 | <0.0250 | <0.0250 | NC | 2 | |

**Table S2.** Caco-2 cell permeability assays for UMK57 (performed by Cyprotex Discovery, Macclesfield, UK).

| **Compound** | **Caco2 Permeability Dynamic (Design1=A2B, Design2=B2A, BufferA/B=HBSS/HBSS, pH_A/B=7.4/7.4** | | | | | | | | | | | | | |
|---|---|---|---|---|---|---|---|---|---|---|---|---|---|---|
| | **Direction=A2B** | | | | | | **Direction=B2A** | | | | | | **Refflux Ratio (Mean $P_{app}$ B2A / Mean $P_{app}$ A2B)** | **Comments** |
| **Name** | **$P_{app}$ ($10^{-6}$ $cms^{-1}$)** Replicate 1 | Replicate 2 | **Mean $P_{app}$ ($10^{-6}$ $cms^{-1}$)** | **SD** | **n** | **Mean % Recovery** | **$P_{app}$ ($10^{-6}$ $cms^{-1}$)** Replicate 1 | Replicate 2 | **Mean $P_{app}$ ($10^{-6}$ $cms^{-1}$)** | **SD** | **n** | **Mean % Recovery** | | |
| antipyrine | 42 | 40,1 | 41,1 | 1,31 | 2 | 91,7 | 40,9 | 42,2 | 41,6 | 0,909 | 2 | 93,2 | 1,01 | Human absorption = 97% |
| atenolol | 0,295 | 0,377 | 0,336 | 0,0584 | 2 | 91,2 | 0,667 | 0,511 | 0,11 | 0,11 | 2 | 98 | 1,75 | Human absorption = 50% |
| talinolol | 0,27 | 0,313 | 0,291 | 0,0308 | 2 | 91,1 | 9,47 | 11,2 | 1,25 | 1,25 | 2 | 90,2 | 35,5 | P-gp substrate |
| estrone 3-sulfate | 0,608 | 0,432 | 0,52 | 0,124 | 2 | 69,4 | 38,4 | 40 | 1,1 | 1,1 | 2 | 82,3 | 75,3 | BCRP substrate |
| **UMK57** | 19,2 | 18,4 | 18,8 | 0,529 | 2 | 59,6 | 45,7 | 28,9 | 37,3 | 11,9 | 2 | 59,7 | 1,98 | - |

5

**Table S3.** CYP-mediated metabolic clearance of UMK57 (provided by Cyprotex Discovery, Macclesfield, UK).

| **Compound** | **Metabolic Stability (Species=Human, HasQCs=No)** | | | | |
|---|---|---|---|---|---|
| **Name** | **$CL_{int}$ (µL/min/mg protein)** | **SE $CL_{int}$** | **$t_{1/2}$ (min)** | **n** | **Comments** |
| dextromethorphan | 45,5 | 3,68 | 30,5 | 5 | - |
| verapamil | 276 | 15,6 | 5,02 | 3 | 30 and 45 minute time points excluded. |
| UMK57 | 257 | 7,61 | 5,38 | 4 | Minus cofactor control low (55% of 0min). Possible chemical instability or non-cofactor dependent enzymatic degradation. 45 minute time point excluded. |

| **Compound** | **Metabolic Stability (Species=Mouse, HasQCs=No)** | | | | |
|---|---|---|---|---|---|
| **Name** | **$CL_{int}$ (µL/min/mg protein)** | **SE $CL_{int}$** | **$t_{1/2}$ (min)** | **n** | **Comments** |
| diazepam | 688 | 15,5 | 2,02 | 3 | 30 and 45 minute time points excluded. |
| diphenhydramine | 70,1 | 4,03 | 19,8 | 5 | - |
| UMK57 | 922 | 0 | 1,5 | 2 | $CL_{int}$ calculated from 0 and 5 minute time points only. 15, 30 and 45 minute time points excluded. |

**Table S4.** Fibroblasts from skin biopsies of Caucasian males used in this study. n/a – not applicable; DA – Day; YR – Years; MO – Months.

| ***Catalog ID*** | ***Description*** | ***Collection*** | ***Age*** | ***Sex*** | ***Tissue Type*** |
|---|---|---|---|---|---|
| DFM021711A | Apparently healthy individual | Zen Bio | 1 DA | Male | Foreskin |
| GM21811 | Multiple cell types from same subject - foreskin Apparently healthy individual | NIGMS Human Genetic Cell Repository | 1 DA | Male | Foreskin |
| GM05659 | Apparently healthy individual | NIGMS Human Genetic Cell Repository | 1 YR | Male | Skin, Chest |
| GM08398 | Apparently healthy individual | NIGMS Human Genetic Cell Repository | 8 YR | Male | Skin, Inguinal area |
| GM01651 | Apparently healthy individual | NIGMS Human Genetic Cell Repository | 13 YR | Female | Skin, Arm |
| GM23973 | Apparently healthy individual | NIGMS Human Genetic Cell Repository | 19 YR | Male | Skin, Arm |
| GM08399 | Apparently healthy individual | NIGMS Human Genetic Cell Repository | 19 YR | Female | Skin, Arm |
| GM23964 | Apparently healthy individual | NIGMS Human Genetic Cell Repository | 21 YR | Male | Skin, Arm |
| GM23976 | Apparently healthy individual | NIGMS Human Genetic Cell Repository | 22 YR | Male | Skin, Arm |
| HGADFN003 | Clinically affected by Hutchinson-Gilford Progeria. Heterozygotic mutation on LMNA Exon 11. | PRF Cell and Tissue Bank | 2 YR 0 MO | Male | n/a |
| HGADFN167 | Clinically affected by Hutchinson-Gilford Progeria. Heterozygotic mutation on LMNA Exon 11. | PRF Cell and Tissue Bank | 8 YR 5 MO | Male | n/a |
| HGADFN169 | Clinically affected by Hutchinson-Gilford Progeria. Heterozygotic mutation on LMNA Exon 11. | PRF Cell and Tissue Bank | 8 YR 6 MO | Male | n/a |
| AG05397 | Trisomy 21 47, XY+21 | NIA Aging Cell Culture Repository | 1 YR | Male | Skin, thorax/abdomen |

| | | | | | |
|---|---|---|---|---|---|
| AG06922 | Trisomy 21<br>47, XY+21 | NIA Aging Cell Culture Repository | 2 YR | Male | Skin, thorax/abdomen |
| AG04823 | Trisomy 21<br>47, XY+21 | NIA Aging Cell Culture Repository | 5 YR | Male | Skin, thorax |
| GM02767 | Trisomy 21<br>47, XY+21 | NIGMS Human Genetic Cell Repository | 14 YR | Female | Skin, unspecified |
| AG08941 | Trisomy 21<br>47, XY+21 | NIA Aging Cell Culture Repository | 19 YR | Female | Skin, arm |
| AG08942 | Trisomy 21<br>47, XY+21 | NIA Aging Cell Culture Repository | 21 YR | Male | Skin, arm |
| GM04928 | Trisomy 21<br>47, XY+21 | NIGMS Human Genetic Cell Repository | 27 YR | Male | Skin, thorax |

**Table S5.** List of primers used for RT-qPCR analysis of human genes.

| Gene Symbol | Forward primer (5' to 3') | Reverse primer (5' to 3') |
|---|---|---|
| *KIF2C* | CTCAGTTCGGAGGAAATCATGTC | TGCTCTTCGATAGAATCAGTCAC |
| *HPRT1* | TGACCAGTCAACAGGGGACA | CTGCATTGTTTTGCCAGTGTCA |
| *TBP* | GAGCCAAGAGTGAAGAACAGTC | GCTCCCCACCATATTCTGAATCT |
| *XRCC5* | CCAGCTTTGAGGAAGCGAGT | AGGCTCGGATGCAGTCTATG |
| *XRCC6* | AAGCCGTTGGTACTGCTGAA | CAGGGTTGAGCTCCCAATCA |
| *PARP1* | GGAGTGGATGAAGTGGCGAA | ATCAGGTCGTTCTGAGCCTTT |
| *RAD50* | GCGGGAAAGACGACCATCAT | TGAGCAACCTTGGGATCGTG |

**Table S6.** List of primers used for RT-qPCR analysis of mouse genes.

| Gene Symbol | Forward primer (5' to 3') | Reverse primer (5' to 3') |
|---|---|---|
| *Kif2C* | GCTTAATTCACCCCGCCAATAT | ATCGTCAATGTCAATCTCTTT |
| *Il6* | GGATACCACTCCCAACAGACC | TGCCATTGCACAACTCTTTTCT |
| *Mmp1* | GGAGCCCTGATGTTTCCCAT | GTCTTCATCGCCTGGACCATA |
| *Mmp13* | AGCAGTTCCAAAGGCTACAACT | ATGGGAAACATCAGGGCTCC |
| *Cdkn2a* | CGTACCCCGATTCAGGTGATG | AGAAGGTAGTGGGGTCCTCG |
| *Cdkn1a* | TCTTGCACTCTGGTGTCTGAG | GCTTGGAGTGATAGAAATCTGTC |
| *Serpine1* | AGAAAGACCGCAAGAACCCG | TCCAACTCGCCTTATTCCTTCT |
| *Hprt1* | CAGTCCCAGCGTCGTGATTA | CACTTTTTCCAAATCCTCGGCA |
| *Gapdh* | AGGTCGGTGTGAACGGATTT | ATGAAGGGGTCGTTGATGGC |
| *Parp1* | GAGTGGAGTACGCGAAGAGC | TCGAACATGGGTGACTGCAC |
| *Rhno1* | CGTCACTTCCTGGGTGTCA | GTAGATCTCCGAGTTGGGCG |
| *Fancd2* | AGTGGCAGAAGACGTTAGTCA | AAGGGGCATGGTTTTGGAGG |
| *Fancb* | TCCCAAAGAAGGAATCAGTAGAACA | AAATTCGGGGCATGGGTAGG |
| *Xrcc6* | TCCGCTTCACATACAGGAGC | CTACCACTTGCTCCGACTCC |
| *Rad51* | ACCAACCAGGTAGTAGCCCA | GGTACAGCCTGGTGGTTGAC |
| *Foxm1* | CTGTGAGGGTCAAAGCTTGC | GCAGCCTCCGTCTTTTGAGA |
